# Orientation-tuned surround suppression exhibits a unique laminar signature in human primary visual cortex

**DOI:** 10.64898/2025.12.12.694066

**Authors:** Joseph H. Emerson, Karen Navarro, Cheryl A. Olman

## Abstract

Spatial context modifies visual perception by enhancing novel and salient features over spatially redundant features in the underlying neural code of primary visual cortex (V1). Although multiple intracortical pathways contribute to contextual modulation, their specific contributions to different types of contextual modulation are not fully understood. Leveraging the distinct laminar connectivity patterns of feedforward, feedback, and lateral pathways, we used ultra-high-resolution fMRI (7T T_2_*-weighted, 0.6 mm isotropic resolution) to infer their relative contributions to contextual modulation in V1 by analyzing blood-oxygenation-level-dependent (BOLD) signal across cortical depth. Participants viewed sine-wave grating disks embedded in large surround gratings. Segmentation cues were introduced or removed by manipulating the relative phase and orientation of the surround gratings, yielding three contextual conditions and a surround-only condition to measure the effects of context in the absence of feedforward input. Our analysis isolated the effects of orientation-tuned surround suppression (OTSS) from orientation-independent border-induced modulation (BIM). The results show that BOLD laminar profiles differ by modulation type: OTSS was absent from deep layers, whereas BIM was more broadly distributed. We also find that voxels at all depths are driven by spatial context in the absence of feedforward input, which accords with the discovery of contextually-driven neural responses in mammalian V1. These laminar differences likely reflect different proportional contributions of feedback from higher-order visual areas and long-range lateral connections within V1. Our findings help to explicate the contributions of recurrent processing to visual contextual modulation and its impacts on laminar-dependent BOLD fMRI.

**Significance Statement:** Our sensory experience requires interpretation: the brain uses both hyper-local and large-scale scene cues to process sensory inputs. In primary visual cortex (V1), this contextual modulation arises from a mixture of intra- and inter-regional neural connections. Because these computations occur at different depths in the cortical gray matter, sub-millimeter resolution fMRI offers an opportunity to quantify the separate contributions of these neural pathways. Using a visual surround suppression paradigm, we measured a distinct depth profile for intra-regional contextual modulation, separating this orientation-tuned signal from other forms of border-induced modulation. This work validates an important new tool for understanding both typical and atypical neural network architectures in the human brain.

## Introduction

Primary visual cortex (V1) acts not as a passive relay but as an active processor, in which neural responses to local visual features are shaped by the broader spatial context. Contextual modulation plays a central role in adaptive visual perception, emphasizing salient features and minimizing redundancy in the neural code (Coen-Cagli, Kohn, & Schwartz, 2015; Li, 2002). While some contextual effects are inherited from subcortical regions (Ozeki et al., 2004), contextual modulation in V1 also relies on recurrent connections, including long-range lateral projections within V1 (Lamme & Roelfsema, 2000; Adesnik, Bruns, Taniguchi, Huang, & Scanziani, 2012; Shushruth et al., 2012; Sato, Häusser, & Carandini, 2014), and feedback from higher-order visual areas (HVAs) (Lamme & Roelfsema, 2000; Bair, Cavanaugh, & Movshon, 2003; Nassi, Lomber, & Born, 2013; Nurminen, Merlin, Bijanzadeh, Federer, & Angelucci, 2018; Vangeneugden et al., 2019), which contribute to integration across spatial scales and feature types (Angelucci et al., 2017, 2002; Kirchberger et al., 2021).

In parafoveal and peripheral regions of the visual field, the presence of a visual stimulus surrounding a V1 neuron’s classical receptive field (CRF) typically suppresses responses to stimuli within the CRF (Angelucci et al., 2017; Hubel & Wiesel, 1968). Surround suppression of neural responses is known to correlate with behavioral measures of perception (Zenger-Landolt & Heeger, 2003; Shushruth et al., 2013; Vanegas, Blangero, & Kelly, 2015; Schallmo, Grant, Burton, & Olman, 2016; Self et al., 2016), suggesting that suppression of neural responses may underlie effects of visual context on perceptual judgments of features such as contrast, orientation, and contours. In V1 neurons, surround suppression shows a complex dependence on the orientation of the center and surround stimuli (Shushruth et al., 2012; Sillito, Grieve, Jones, Cudeiro, & Davls, 1995) but is generally strongest when the center and surround are at or near the preferred orientation of a neuron (Cavanaugh, Bair, & Movshon, 2002b; Jones, Wang, & Sillito, 2002; Nothdurft, Gallant, & Van Essen, 1999). Lateral pathways in V1 that preferentially link neurons with similar orientation preferences (Malach, Amir, Harel, & Grinvald, 1993) have been suggested to explain orientation tuned surround suppression (OTSS), an idea that is supported by the discovery of V1-intrinsic microcircuits facilitating OTSS in mice (Adesnik et al., 2012). However, in primates, lateral connections can only support suppression between nearby stimuli, since the spatial extent of these connections is limited (Angelucci & Bullier, 2003). Furthermore, individual cell recordings have shown that OTSS can take effect within 15-25 ms after the stimulus-evoked response onset (Bair et al., 2003; Henry, Joshi, Xing, Shapley, & Hawken, 2013; Knierim & Van Essen, 1992; Nothdurft et al., 1999). Such a short latency requires invoking fast-conducting myelinated axons from HVAs (Angelucci & Bullier, 2003; Bair et al., 2003; Girard, Hupé, & Bullier, 2001). Silencing HVAs reduces the amount of surround suppression in V1 in awake mice (Vangeneugden et al., 2019) and macaques (Nassi et al., 2013; Nurminen et al., 2018), supporting a role for HVAs in OTSS. Still, significant silencing of HVAs does not completely extinguish surround suppression, which strongly suggests that V1-intrisic and/or subcortical mechanisms also contribute. Therefore, OTSS is likely influenced by a combination of neural pathways that include both V1-intrinsic lateral circuits and feedback modulation by HVAs.

Higher-order scene statistics, such as the presence of borders demarcating the boundaries of objects, also influence contextual modulation in V1. Responses of V1 to foreground regions are enhanced over background regions, a phenomenon called figure-ground modulation (FGM) (Lamme, 1995; Lamme, Rodriguez-Rodriguez, & Spekreijse, 1999; Self et al., 2016; Schnabel et al., 2018). It is unlikely that FGM originates in V1; mechanisms for assigning border ownership, a prerequisite to object detection, are likely articulated by extrastriate regions, including V2 and V4 (Zhou, Friedman, & Von Der Heydt, 2000; Roelfsema, Lamme, Spekreijse, & Bosch, 2002). Consistent with FGM requiring additional extrastriate processing, FGM in primate V1 has an even longer latency than OTSS (*>*30 ms after stimulus-evoked response onset) (Lamme, 1995; Lamme et al., 1999; Lamme & Roelfsema, 2000; Self, van Kerkoerle, Super, & Roelfsema, 2013) and is most pronounced in cortical layers targeted by feedback from HVAs (Self et al., 2013). Furthermore, optogenetic silencing of HVAs in mouse (Kirchberger et al., 2021) and lesioning of HVAs in macaques (Lamme, Supér, & Spekreijse, 1998) reduce or extinguish FGM in V1 demonstrating a causal role for feedback.

Despite this thorough understanding of OTSS and FGM in animal models, there remain gaps in our understanding of these forms of contextual modulation in the human visual system. The present study was designed to validate and expand upon what is known about OTSS and FGM in humans. Contextual modulation manifests in the blood-oxygenation-level-dependent (BOLD) signal of early visual cortex (Poltoratski & Tong, 2020; Tajima et al., 2010; Williams, Singh, & Smith, 2003; Zenger-Landolt & Heeger, 2003), and laminar fMRI provides a new opportunity to study the contributions of feedforward, feedback, and lateral pathways (L. Huber et al., 2017; Koopmans, Barth, & Norris, 2010; Lawrence, Formisano, Muckli, & de Lange, 2019). Using an established contextual modulation paradigm that uses either a phase shift or rotation of a circular grating patch to define a figure against an extended sinusoidal grating surround (Kirchberger et al., 2021; Self et al., 2016), our primary goal was to isolate OTSS from figure-ground segmentation effects. Based on the available evidence for the cortical origins of OTSS and FGM, we hypothesized that OTSS would disproportionately modulate superficial layers.

A notable challenge in studying scene segmentation effects in human V1 (as opposed to anesthetized animals) is that effects of attention are easily confounded with FGM. Object-based attention modulates V1 signals ~160 ms after stimulus-evoked response onset (Poort et al., 2012); in the sluggish fMRI response, rapid border ownership signals, or other feedback processes resulting from grouping or object recognition, are not readily separated from effects of attention allocated to objects defined by the borders. A key comparison in this study is the comparison of V1 surround suppression by parallel surrounds in the presence and absence of a border. To minimize modulation of or by attention, participants in the present study were directed to perform a demanding fixation task. However, covert (and unreported) attention to the disks that were visible in some conditions but not others could contribute to the measured fMRI response whenever stimulus edges were present. Thus, we will use the term border-induced modulation (BIM) throughout to indicate the modulation of the V1 fMRI signal by the presence of an encircling edge in the scene, with the understanding that this comprises the FGM signals of interest as well as potential object-based attention effects. Due to the likely feedback origin of FGM as well as attentional effects in V1, we hypothesized that BIM would primarily modulate both superficial and deep layers.

## Results

Small peripheral sine-wave grating disks (target stimuli) were presented simultaneously with a large surround grating that was either matched or unmatched to the orientation or phase of the target stimuli (Fig. 1A). To ensure that attention was maintained at fixation, the participants performed an orientation discrimination task on a small central grating disk (see Methods). BOLD data covering occipital regions of cortex were collected in a 7T MRI scanner using a T_2_*-weighted acquisition at 0.6 mm isotropic resolution. We analyzed data from 12 participants to quantify the profiles of OTSS and BIM both across the cortical surface and through the cortical depth.

**Figure 1.**
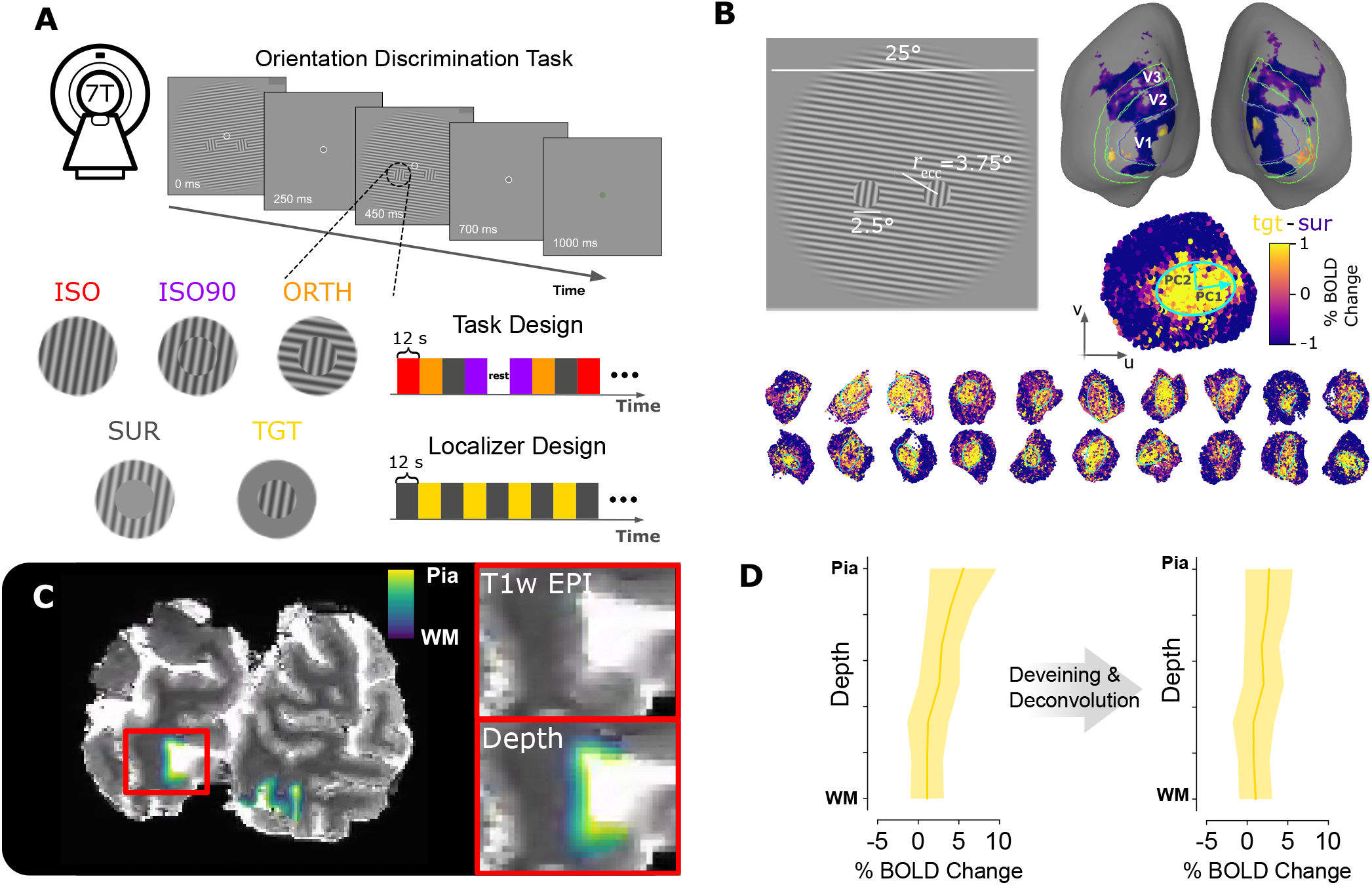
Experimental design and analysis pipeline. (A) Top: A single trial of the foveal orientation-discrimination task that participants performed while undergoing functional scanning at 7T is illustrated. The white circles enclosing the central grating disks are enlarged for visibility. Bottom left: Three task conditions modified the features of the surround relative to the targets and two additional target-only and surround-only conditions localized the spatial representation of the stimulus in V1. Bottom right: Task scans measured the effects of *iso, iso90, orth*, and *sur* conditions on BOLD responses, while a localizer scan identified target- and surround-selective voxels. Conditions for both scans were presented in 12 s blocks of 8 trials each. (B) Left: Measurements are illustrated on an example stimulus. Right: The voxel-wise contrast between *tgt* and *sur* conditions was projected onto a 3D rendering of the gray matter surface for each participant after significance thresholding based on the *tgt* - *sur* contrast (*p <* 0.01 without correction for multiple comparisons). Boundary estimates for cortical regions V1, V2, and V3 were first identified using anatomical landmarks from the Benson atlas (Benson et al., 2014) for each participant. For five of the twelve participants, functional retinotopy scans provided further validation and refinement of these boundaries (V1 - purple outlines, V2 - dark green outlines, V3 - light green outlines) (see Methods). Bottom: Flattened cortical patches were 10mm-radius disks centered on the target-selective regions identified by the *tgt* - *sur* contrast on the surface projections. The patches of the cortical ribbon were then extracted and transformed in volumetric space, yielding new flattened coordinates, *u* and *v* (see Methods). The first and second principal components of the spatial covariance matrix of the flattened coordinates (*u,v*) of the target-selective voxels (yellow) were used to define an elliptical boundary for the target-selective ROIs. Each of the twenty-one ROIs used in subsequent analyses is shown. (C) Left: Voxel depth assignment within the extracted cortical ribbon volume was determined from the in-session T_1_w-EPI. Top right: An enlarged view of the T1w EPI image in the region outlined by the red rectangle on in the left subpanel is shown. Bottom right: The same enlarged T1w EPI is shown with gray matter voxels color-coded according to depth. (D) Left: Average percent BOLD change across depth for a single ROI during the target-only condition. The shaded region indicates standard deviation. Right: Deveining was performed using a vein mask derived from the mean-normalized variance of the residual time series data (see Methods). A GLM-based depth deconvolution approach was then used to remove the pial bias of the resulting ROI depth profiles (Markuerkiaga et al., 2016) (see Methods and Supplementary Materials).

### Localization of target-selective voxels

Regions of interest (ROIs) were defined from an independent localizer that presented the target and surround regions in block alternation. Because different mixtures of feedforward and feedback effects are expected at the center versus the edges of the target region of interest, we took additional steps to define a flattened coordinate system that would allow us to isolate responses to different regions of the target (see Methods for detail). Due to cortical magnification, the target-selective voxels formed oblong regions on the cortical surface. In brief, we defined a distance coordinate, *σ*, as one standard deviation of the spatial distribution of target-selective voxels along each principal component of the patch. This characterized an ellipse that conformed to the oblong shape of the target-selective patch (Fig. 1C).

Fig. 2B shows the variation in voxel preference for target versus surround stimulus as a function of distance from the patch center. Due to how we defined *σ*, most voxels responding more strongly to the target are located within a 2*σ* ring; the retinotopic location of the edge between the target and surround is between 2*σ* and 3*σ*. To ensure that we quantified OTSS and BIM in voxels with population receptive fields (pRFs) that were well localized to the target, we chose an ROI boundary of 1*σ* (1.53 ± 0.36 mm along the minor axis). The area within 1*σ* of the ROI center (11.93*±*4.97 mm^2^) conforms to the theoretical scale of the spatial representation of the central 1° of the target disk on the cortical surface according to the standard cortical magnification for human V1 at 3.75° eccentricity (Fig. 2D-E, see Methods). The flat *tgt-sur* contrast within 1*σ* confirms that voxels within this boundary were driven only by the target region and did not have PRFs that overlapped with the target edge. Voxels within 1*σ* were significantly target-selective (*p <* 0.05) in all depth bins used in subsequent analysis except the deepest of the seven bins, at the gray matter/white matter boundary (Fig. 2C & Table S6).

**Figure 2.**
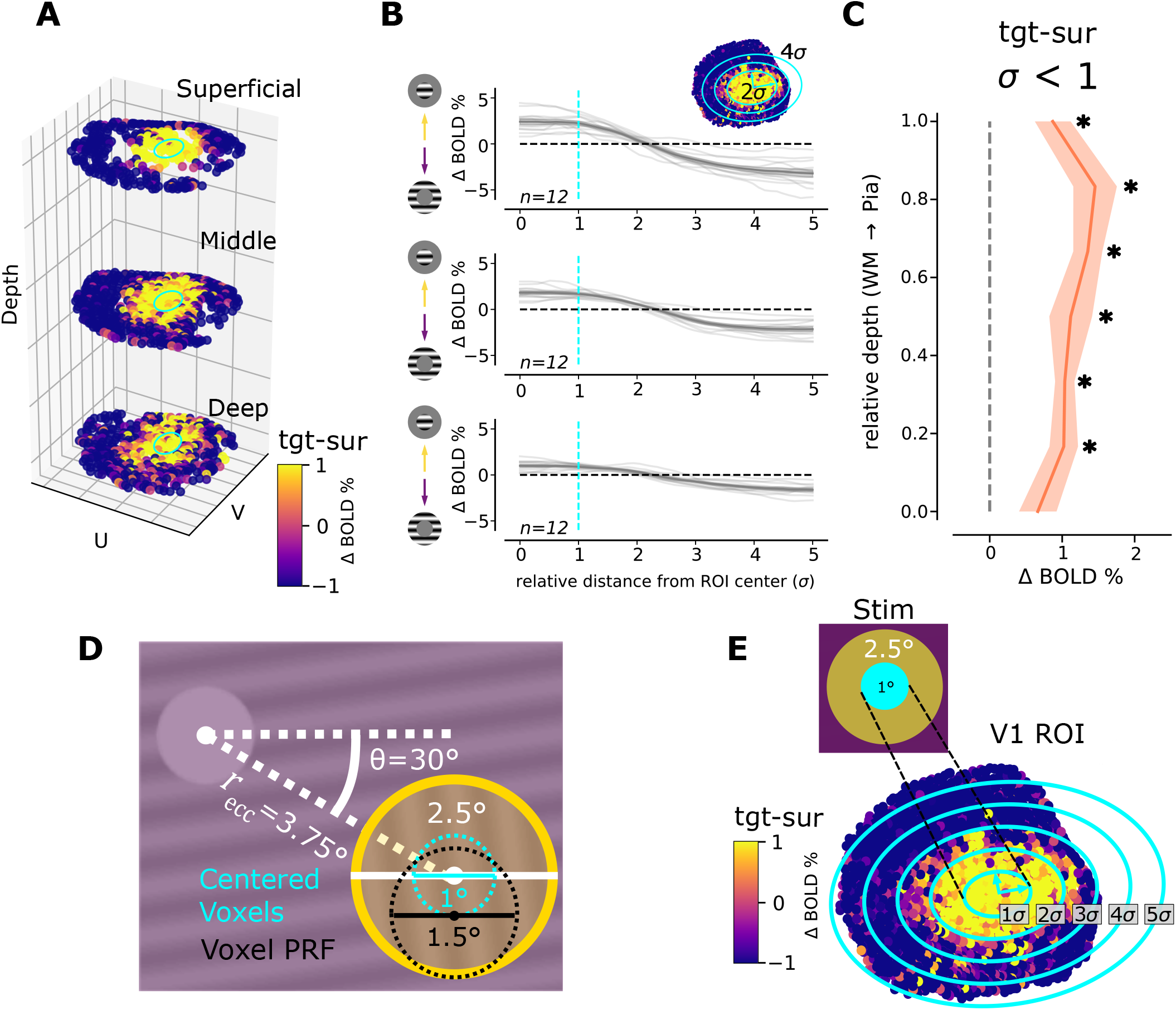
Assessment of ROI localization. (A) Voxel centroids are plotted at three depth bins on a flattened coordinate system to illustrate the target- and surround-selective regions. (B) Radial profiles for each ROI were constructed to assess the localization of target-selective voxels. For each participant, smoothed *tgt* - *sur* profiles are shown at each of three depth bins (light lines). The average across participants is shown in bold. Positive values indicate stronger responses to the target-only condition while negative values indicate stronger responses to the surround-only condition. The inset demonstrates how the radial dimension of each ROI is quantified, with *σ* denoting the base scaling of the ellipse. (C) The grand average *tgt* - *sur* contrast within a radius of 1*σ* is plotted at seven depth bins. Stars indicate that the contrast at the corresponding depth bin is significantly greater than zero (*p <* 0.05, single-sample two-sided permutation test, *N*_*perm*_ = 4096, FDR-corrected, see Table S6). All shaded regions indicate standard error of the mean. (D) Dimensions of the stimulus targets are shown. The black dotted circle indicates the expected maximum pRF size for voxels centered at 3.75° eccentricity. Given this pRF size, pRFs centered within the cyan dotted circle do not overlap with the edge of the target. (E) A miniaturized representation of the target region appears in yellow with the cyan circle indicating the spatial region containing pRFs that do not overlap with the target edge. According to the standard cortical magnification of visual space in V1, the cyan region is represented by approximately the area within 1*σ* of the ROI center (11.93*±*4.97 mm^2^).

### Localization of context modulation on the cortical surface

Selective pairwise comparisons of three stimulus conditions presented in block-design task scans (Fig. 1A) isolated the effects of OTSS from BIM. In the *iso* condition, we matched the orientation and phase of the surround to the target gratings, such that the targets were indistinguishable from the surround (Fig. 3A). In the *iso90* condition, the phases of the target and surround gratings were offset by 90°, creating a luminance boundary between the target and surround (Fig. 3B). This boundary perceptually segregated the target regions from the surround gratings and enhanced the responses of target-selective voxels relative to the *iso* condition. Thus, BIM was operationalized as a contrast between *iso90* and *iso* conditions. In a third condition called *orth*, the surround stimulus was rotated 90° relative to the target stimuli (Fig. 3C). Because both *iso90* and *orth* contained edges, the contrast between these two can isolate OTSS.

**Figure 3.**
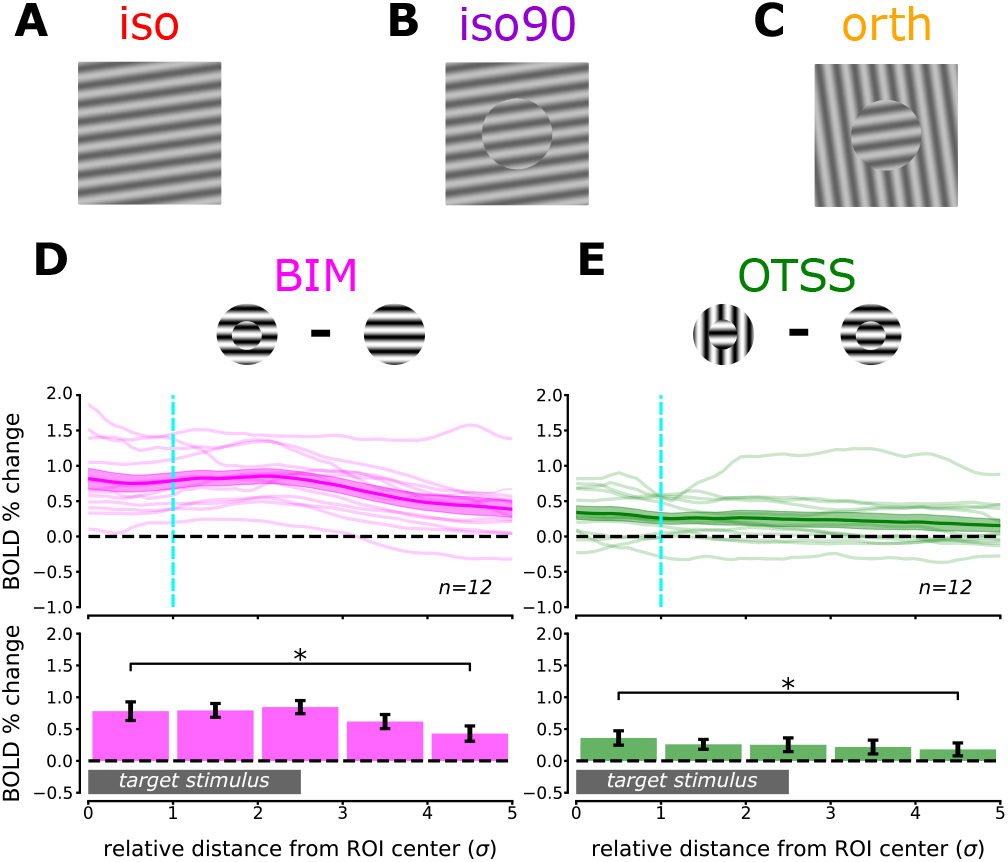
Contextual modulation along the cortical surface. (A-C) Example stimuli for the *iso* (A), *iso90* (B), and *orth* (C) conditions are shown. (D-E) BIM (*iso90* - *iso*) (D) and OTSS contrasts (*orth - iso90*) (E) are plotted along the radial axis of the ROI. *Top*: Individual participant profiles are indicated by light-colored lines. Bold lines indicate grand average profiles over all participants. The vertical cyan line indicates the boundary at which we estimate voxel pRFs to be well localized to the target. *Bottom*: Bar plots show the grand average at discrete radial bins. The gray bar indicates the approximate size of the V1 representation of the target stimulus. Stars with brackets indicate significant differences (*p <* 0.05) between 0-1*σ* (our target ROI) and 4-5*σ* (where voxel pRFs are localized to the surround) (BIM: *p* = 0.010 and OTSS: *p* = 0.013, paired two-sided permutation tests, *N*_*perm*_ = 4096, FDR-corrected, see Tables S16 & S17). Shaded regions and error bars in all plots indicate standard errors of the mean.

Both forms of contextual modulation measured in the BOLD signal were spatially selective. We observed a significantly greater release from surround suppression in voxels with pRFs that were well localized to the center of the target region (*r <* 1*σ*) than in voxels with pRFs well outside the target region (4*σ ≤ r <* 5*σ*) (BIM: mean difference= 0.352, *p* = 0.010, Fig. 3D; OTSS: mean difference= 0.179, *p* = 0.013, Fig. 3E; paired two-sided permutation test, *N*_*perm*_ = 4096, FDR-corrected, see Tables S16 & S17).

In addition to showing strong BIM at the center of the target, the grand average radial BIM profile shows a slight peak between 2*σ* and 3*σ*, where the edge of the target grating is located. Since the *iso90* condition contains edge-induced contrasts that are absent in the *iso* condition, this is likely due to feedforward responses in neurons with high spatial frequency selectivity and receptive fields that overlap the edge. However, this feedforward edge response is spatially limited, as demonstrated by the rapid decrease in the 3*σ*-4*σ* and 4*σ*-5*σ* bins. The enhanced responses to the *iso90* condition within the center of the ROI (*r ≤* 1*σ*) relative to the distal regions of the ROI (4*σ < r ≤* 5*σ*) is indicative of preferential responses to the target over the surround during the *iso90* condition independent of any edge-induced feedforward contributions.

Since both the *iso90* and the *orth* conditions contained edge-induced contrasts, the OTSS contrast isolates signals related to the orientation differences between the target and surround. Consequently, we observed no peak in the radial profile of OTSS near the retinotopic representation of the edge (2*σ < r ≤* 3*σ*, Fig. 3E).

### Distinct Laminar Signatures of BIM and OTSS in Human V1

Consistent with the prediction that condition contrasts isolating BIM (*iso90* - *iso*) would reveal the strongest effects in superficial and deep layers (the primary targets of feedback projections from HVAs), we observed peaks in superficial and deep bins in the average BIM depth profile (Fig. 4D). However, neither Kruskal-Wallis tests nor one-way ANOVAs indicated a main effect of depth in the BIM profile (Kruskal-Wallis *H* = 5.15, *p* = 0.525; one-way ANOVA *F* = 0.922, *p* = 0.484, see Tables S1, S2), so we cannot conclude that modulation was absent in middle layers.

**Figure 4.**
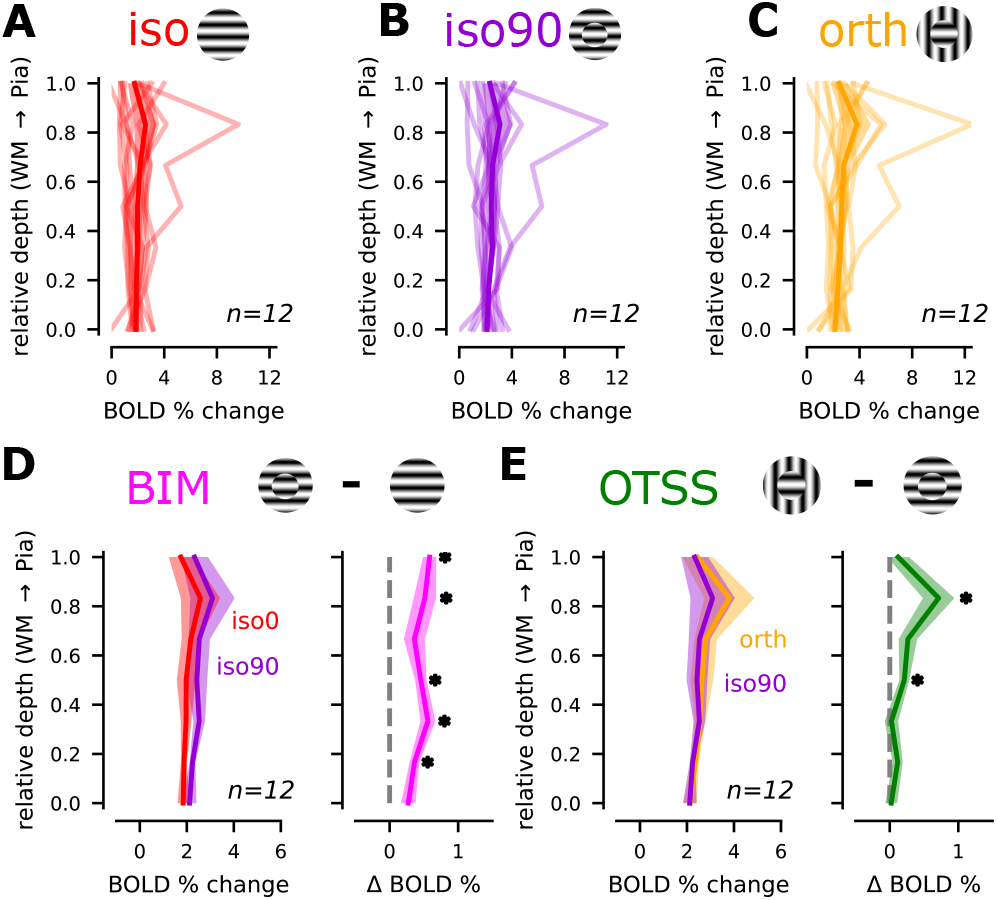
Contextual modulation of BOLD laminar profiles. (A-C) Depth profiles for the *iso* (A), *iso90* (B), and *orth* (C) conditions for the target-selective ROIs are shown for individual participants (light-colored lines) and grand averages across participants (bold lines). (D) Left: Grand average depth profiles of target-selective ROIs are plotted for the *iso90* (purple) and *iso* (red) conditions. Right: The grand average depth profile for the *iso90* - *iso* contrast (BIM) shows the average across all twelve participant profiles. (E) Left: Grand average profiles of target selective voxels are plotted for the *orth* condition (orange) and the *iso90* condition (purple). Right: The grand average profile for the *orth*-*iso90* contrast (OTSS). Shaded regions in all panels indicate standard error of the mean across participants. Stars indicate significant release from suppression (*p <* 0.05).

Given that long-range lateral connections likely contribute to OTSS (Kirchberger et al., 2021; Adesnik et al., 2012) in addition to feedback (Angelucci & Bullier, 2003; Bair et al., 2003; Girard et al., 2001; Nassi et al., 2013; Nurminen et al., 2018; Vangeneugden et al., 2019), we predicted that the strongest impacts of OTSS (*orth* - *iso90*) would be observed in superficial layers, where long-range lateral connections in V1 are particularly dense (Rockland & Pandya, 1979). The laminar profiles of the OTSS contrast indeed revealed a main effect of depth (Kruskal-Wallis *H* = 12.8, *p* = 0.046; one-way ANOVA *F* = 4.11, *p* = 0.011, see Tables S3 and S4). Significant effects were measured in middle to superficial bins (single-sample two-sided permutation test, *N*_*perm*_ = 4096, FDR-corrected, see Table S8), with a clear superficial peak (Fig. 4E). The restriction of OTSS to superficial and middle bins is consistent with population-level electrophysiological recordings from macaques (Henry et al., 2013; Self et al., 2013) and mice (Self et al., 2014) showing the most prominent orientation-tuned suppression in supragranular layers and upper layer 4.

While significant main effects of depth (two-way repeated measures ANOVA *F* = 5.74, *p* = 7.41 *×* 10^−5^) and condition contrast (two-way repeated measures ANOVA *F* = 6.94, *p* = 0.027) were found, no significant interaction between depth and condition contrast was observed (two-way repeated measures ANOVA *F* = 1.53, *p* = 0.183, see Table S5). Therefore, while the OTSS profile lacked deep bin effects observed in the BIM profile, differences were not statistically significant in this sample. Nevertheless, the independent BIM and OTSS profiles are highly suggestive of laminar activity patterns predicted by the structural distribution of lateral and feedback connections respectively, as previously discussed.

We noted the presence of a significant outlier participant with large percent BOLD change in all conditions with respect to the rest condition. This participant met all inclusion criteria and the large responses did not appear to be due to any signal artifacts. We repeated statistical analyses without this participant and found that the trends observed across participants were not disproportionately driven by this outlier (see Fig. S5 and Tables S18-S25).

### BOLD response in V1 is driven by the spatial surround in the absence of feedforward input

Despite conservative localization of target-selective regions (Fig. 2A), we found unexpected, strong modulation of BOLD responses in target-selective regions during the surround-only (*sur*) condition (Fig. 5A). While response magnitudes for *iso* and *sur* were matched in the center of the target regions (Fig 5C), the *sur* condition drove stronger responses near the boundary and at more distal radii (Fig. 5D). The increased response near the boundary and in distal regions may reflect the impact of edge-induced contrasts and/or broad amplification of V1 responses induced by the presence of the gray disks.

**Figure 5.**
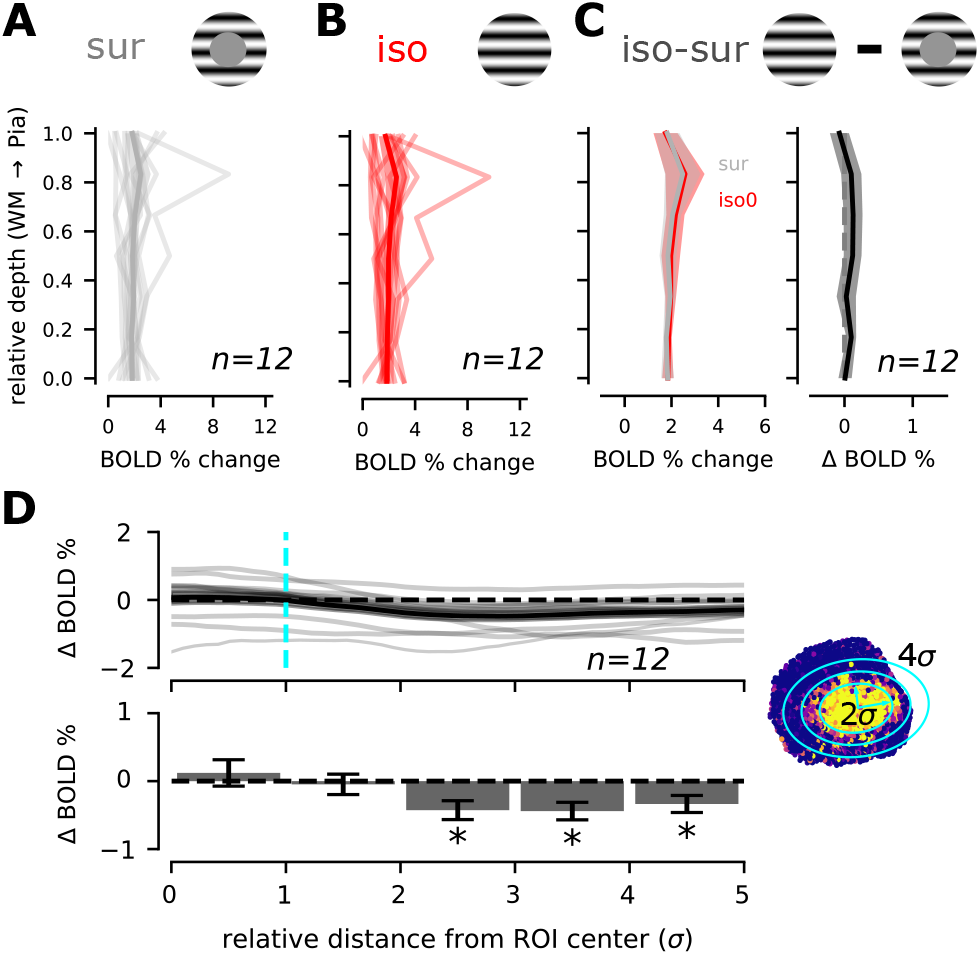
BOLD response in the absence of feedforward input. (A-B) Depth profiles for the *sur* (A) and *iso* (B) conditions are shown for individual participants (light-colored lines) and grand averages across participants (bold lines). (C) Left: Direct comparison of the surround-only and *iso* profiles across depth reveal no significant differences (*p <* 0.05, single-sample two-sided permutation test, *N*_*perm*_ = 4096, FDR-corrected, see Table S9). (D) Top: Individual participant radial *iso-sur* profiles are indicated by light-colored lines. Bold lines show grand average profiles over all participants. The vertical cyan line indicates the boundary at which we estimate voxel pRFs to be well localized to the target. Right: An example ROI indicates the scale of the radial distance; color as in Fig. 2. Bottom: Bar plots show the grand average at discrete radial bins. Stars indicate whether the average was significantly different than zero (*p <* 0.05, single-sample two-sided permutation test, *N*_*perm*_ = 4096, FDR-corrected, see Table S15). All shaded regions indicate standard error of the mean.

A decoding analysis revealed that the surround orientation could be decoded with above-chance accuracy from voxels within 0 *−* 1*σ* during the *sur* condition. However, we were unable to identify a consistent statistical interaction between depth and radial position (Tables S26-S27). Therefore, we were unable to reliably infer the origin of the context-dependent signal in central regions of the ROI (see Supplementary Materials for discussion).

## Discussion

In this study, we isolated the depth-dependent BOLD profiles for orientation-tuned surround suppression (OTSS) from orientation-independent effects caused by the presence of a boundary, which we term border-induced modulation (BIM). We found that OTSS and BIM have unique laminar signatures in human V1 providing further evidence that OTSS is supported by mechanisms distinct from BIM (Self et al., 2016). We furthermore showed that BOLD responses in V1 are not only modulated by spatial context, but that context can drive responses in the absence of feedforward input, a finding consistent with the discovery of feedback receptive fields in non-human animals (Keller, Roth, & Scanziani, 2020; Kirchberger, Mukherjee, Self, & Roelfsema, 2023; Lee, Mumford, Romero, & Lamme, 1998). Our results are in broad agreement with electrophysiological measurements of population activity across cortical laminae, suggesting a close registration of BOLD laminar responses to other population measures of neural activity. These results reveal that both feedback and lateral connections exert a pronounced influence on the BOLD signal, reflecting the impact of spatial context on local representations in V1.

While the mechanisms contributing to the BIM profile were likely complex, the BOLD laminar profile for BIM was consistent with a largely feedback origin of the contextual modulation, peaking in superficial and deep bins where feedback projections primarily terminate (Fig. 4D). The observed trends align with laminar profiles reported in macaque V1 under similar stimulus conditions (Self et al., 2013), though we note that the absence of significant depth-dependent results in the omnibus tests limits the strength of conclusions based on the laminar profile shape alone. BIM was not entirely absent in middle bins, which may be explained by limitations in the spatial resolution of laminar fMRI or by neural mechanisms – both the interactions between layers within V1 as well as mechanisms routing through thalamocortical loops. Indeed, layer 4 is known to display some contextual modulation in V1 (Self et al., 2013, 2014; Kirchberger et al., 2023) and this may be partly explained by similar effects mediated through feedback to the lateral geniculate nucleus (LGN) (Poltoratski, Maier, Newton, & Tong, 2019; Lankow & Usrey, 2022).

Consistent with substantial evidence for orientation-tuning of surround suppression both in V1 neural activity (Cavanaugh, Bair, & Movshon, 2002a; Cavanaugh et al., 2002b; Jones et al., 2002; Nothdurft et al., 1999) and in corresponding behavioral measures of perception (Xing & Heeger, 2000; Vanegas et al., 2015), we observed OTSS that was stronger within the target-selective ROIs than in in adjacent surround-selective regions (Fig. 3B). The asymmetry of OTSS across the boundary is most simply explained by asymmetry in the surround fields: for neurons with CRFs on the surround (convex) side of the curved boundary, less than half the extraclassical receptive field contains orientation contrast, so the release from suppression should be small compared to neurons with CRFs on the concave side of the boundary. However, it is also possible that non-linear interactions between FGM and OTSS rendered residual imprints of FGM on the *orth* - *iso90* contrast. The effects of OTSS were restricted to superficial and middle depths (Fig. 4H) and were remarkably consistent with electrophysiological results from macaque (Henry et al., 2013; Self et al., 2013) and mouse (Self et al., 2014) showing the strongest effects of OTSS in supragranular layers and upper layer 4. Caution is always warranted when interpreting BOLD signals in superficial layers due to the increased susceptibility for the BOLD signal in superficial layers to be contaminated by large pial veins (L. Huber, Uludağ, & Möller, 2019) and the superficial bias of BOLD signal driven by draining vessels (Markuerkiaga et al., 2016). However, we mitigated these effects using depth deconvolution and vein masking. Therefore, the observed superficial layer results likely reflect the underlying local neural activity rather than signals from deeper layers.

Significant BOLD responses in the conservative target ROI during the *sur* condition suggest that V1 is not only modulated but also driven by spatial context. This finding is significant as it accords with recent work in mice (Keller et al., 2020; Kirchberger et al., 2023) and longstanding observations of feedback-driven responses in regions of V1 lacking feedforward drive in macaque (Lee et al., 1998). Interestingly, while previous fMRI work has identified contextual information only in superficial layers of V1 in the absence of feedforward input (Muckli et al., 2015), we find evidence that context drives responses across all layers, which is more consistent with existing electrophysiological evidence.

Another recent study using ultra-high-resolution line-scanning fMRI confirmed that a large annular stimulus positioned well outside the pRFs of voxels in V1 could elicit significant responses in superficial and deep layers, but not middle layers (Heij et al., 2025). They furthermore report that a medium sized annular stimulus that did not directly stimulate the pRFs of the voxels elicited a more complex laminar profile with superficial and deep peaks that shifted toward middle layers. They suggested that the laminar profile for the medium annulus could be caused by a combination of feedback and lateral mechanisms. Their medium-sized stimulus was better matched to the scale and eccentricity of our stimulus, therefore, our inability to measure laminar-dependent effects of the *sur* condition may be caused by our lower spatial resolution relative to Heij et al. (Heij et al., 2025), highlighting the need for improved spatial resolution to appropriately characterize BOLD laminar profiles. Our results in combination with Heij et al. (Heij et al., 2025) are likely to be indicative of a feedback receptive field (Keller et al., 2020) which aids in integrating contextual information across broader spatial regions by incorporating information from HVAs. As pointed out by Kirchberger et al. (Kirchberger et al., 2023), contextually-evoked responses may be described as either a form of FGM whereby the gray disks are perceived as the foreground or, alternatively, as a perceptual filling-in procedure whereby blank spots in the visual field are filled in by the surround. Behavioral measures of perception should be leveraged to provide insights into the perceptual consequences of contextually-driven responses in V1.

Separate analysis of feedforward and feedback effects, in this study, required isolating feedback modulation targeted at figure regions from feedforward drive at border regions. Data were collected in parafoveal regions of V1, where the edges defining the BIM contrast and the *iso*-*sur* contrast would provide strong drive to V1 neurons with preference for high spatial frequencies, which were otherwise not stimulated by the 1.6 cpd sinusoidal gratings used in the study. Due to the duration of scans and the relatively low contrast-to-noise ratio of imaging with 0.6 mm resolution, we found in-session pRF mapping runs to be infeasible. Thus, we rely on the following facts to support the interpretation that responses in our target ROIs were not driven by signal from the border between the target and the surround: image resolution, as estimated by the spatial autocorrelation of residuals after GLM analysis was 0.58 *±* 0.04 mm, and our target ROIs were separated from the retinotopic boundary by a minimum of 1.53 *±* 0.36 mm; while large veins can displace 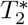-weighted signal several millimeters across cortex, estimates for parenchymal blurring of the 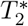-weighted fMRI signal (~1.5 mm) are also smaller than the separation between the stimulus boundary (2*σ* radius) and the conservative target ROIs (1*σ*) used in this analysis (from which voxels with the signature high mean-normalized variance of large veins were removed); radial profiles of the BIM and *iso*-*sur* stimulus contrasts (the two contrasts that differed in the presence of an edge) showed the expected signal increase at the retinotopic location of the edge but were flat inside the *<* 1*σ* ROIs used to assess laminar profiles. Thus, by restricting our analyses to a very small region of interest in the center of the figures, we are able to report laminar profiles of BOLD contrasts that are dominantly driven by feedback, even if complete isolation of signals is not guaranteed.

Although the present study provides evidence for unique laminar profiles for OTSS and BIM in human V1, several limitations – in addition to the question of spatial separation of responses to edges and responses to objects, addressed above – that warrant consideration. First, despite using vein masking and depth deconvolution, residual vascular contamination and depth blurring may influence laminar estimates. Future laminar fMRI studies may benefit from alternative acquisition techniques that are less susceptible to signal contamination from large vessels such as spin-echo BOLD (Duong et al., 2003) and vascular-space occupancy (VASO) fMRI (L. Huber et al., 2019). Second, eye-tracking was not used to confirm that fixation was maintained during stimulus presentation. The target size was kept small to maximize large contextual modulation effects. This small target size meant that eye movements had the potential to bring signal from the surround into the target ROIs. However, the clear and reliable appearance of target-preferring patches in V1, as well as the observed peaks at the target boundary in the BIM and *iso*-*sur* radial profiles, are possible only when fixation is maintained during the functional localizer, which strongly suggests that participants meeting inclusion criteria adequately fixated.

Naturally, alternate interpretations and additional questions remain. Additional work is required to investigate the separate mechanisms contributing to BIM by manipulating attention and task. The current orientation-discrimination task suffered from a strong oblique effect which degraded task accuracy and led to variable control of task difficulty across conditions and participants (see Supplementary Materials). Of great interest would be future studies that explicitly manipulate the degree of attention to the target, or analysis of a bistable stimulus or temporal contextual modulation task that resulted in changes in figure-ground segmentation without changes in borders. Also, while the current study limits its analysis to sinusoidal grating stimuli and contrast-induced edges, future studies should look to extend results to more naturalistic conditions including using random texture elements to control for particular edge-induced contrast patterns. Finally, the conclusions regarding lateral versus feedback pathways are inferential, as we lacked the spatial coverage to directly determine interactions between V1 and HVAs. An aim of future research should be to assess functional connectivity between visual regions as a function of cortical depth, which can serve to corroborate or reject interpretations of the cortical origins of the profiles we report here.

The present study represents an important step toward validating the assumptions underlying the interpretation of laminar fMRI. We use an established framework for measuring various contextual influences on V1 BOLD laminar profiles. This provides a direct point of comparison with other modalities including electrophysiology and optical imaging in animal models using the same or similar paradigms. As suggested elsewhere (Merriam, Gulban, & Kay, 2022), direct comparisons such as these are important for understanding neurohemodynamic coupling. This work also provides an important biomarker of contextual modulation in humans. The continued use of such tools offers the potential to more directly link neural activity to perceptual experience in humans which may have applications for studying the neural mechanisms of diseases affecting perception.

## Materials and Methods

### Participants

Sixteen neurotypical adults (10 female, 5 male, and 1 non-binary), aged 21-48, participated in the study. The University of Minnesota’s Institutional Review Board approved all procedures. Participants gave written informed consent under the Declaration of Helsinki and were compensated at $20/hour.

### Apparatus

Stimuli were displayed using an NEC NP4000 projector at 60 Hz. Participants viewed these through a mirror, 15 cm from their eyes and 65 cm from the screen, resulting in an 80 cm path length from the image to their eyes. The projected image covered a 42 cm x 31.5 cm area on the screen.

### Experimental Design

#### Visual Stimuli

Stimuli, generated using PsychoPy (Peirce et al., 2019), were sine-wave gratings of 1.6 cycles/degree at 40% contrast. The display was divided into three subregions: 1) a circular surround with a diameter subtending 25° at fixation, 2) two target circles subtending 2.5°, each at 3.75° eccentricity and rotated 30° below the horizontal meridian, and 3) a central grating subtending 1.25° (Fig. 1B).

Stimuli were presented in blocks of 8 two-alternative forced-choice (2AFC) orientation discrimination trials, during which the average orientation of the surround and central gratings was one of eight possible orientations arranged in equal intervals between 0° and 157.5°. Each 1.2 s trial had the following sequence: 1) a simultaneous 250 ms presentation of surround, target, and center stimuli, 2) a 200 ms gray screen, 3) another 250 ms presentation of the same stimulus configuration, but with all gratings uniformly rotated slightly in orientation either clockwise or counterclockwise, and 4) a subsequent 300 ms gray screen. Orientation shifts were regulated by a three-down, one-up staircase method (Taylor & Creelman, 1967), with shift values ranging from 1° to 16°. Upon the presentation of the orientation-shifted stimulus, participants had 500 ms to indicate whether the shift was clockwise or counterclockwise. For the final 200 ms, feedback was provided by a change in the fixation circle’s color: green for correct and red for incorrect answers (Fig. 1A). Trials were interspersed with jitters of 0, 300, or 600 ms. Within each block of 8 trials, the average stimulus orientation remained constant. Experiments balanced orientations across all blocks such that every condition (*sur, iso, iso90, orth, tgt*) received the same total number trials for each orientation.

The task component of this experiment was included to ensure that participants remained engaged during the scanning sessions, maintained fixation, and did not attend to the target circles. Due to the short stimulus presentations and response periods, participants generally found the task to be quite difficult. In addition, participants found the task to be substantially more difficult for oblique orientations than cardinal orientations (Fig. S7). Since we did not anticipate the magnitude of the performance difference between cardinal and oblique orientations, separate staircases were not used to independently control the orientation shifts for oblique and cardinal orientations, and the task was more difficult for oblique than cardinal orientations. Therefore, we were not able to robustly estimate orientation discrimination thresholds for eight of the sixteen subjects, but we were able to confirm that participants remained engaged in the task and therefore distracted from the target circles. Due to these methodological difficulties, we did not find participant performance on the task to be relevant to our analysis of contextual modulation effects. See the Supplementary Materials for further discussion of the behavioral data.

#### Experimental conditions

The experiment had two block-design scan types: task scans and localizer scans. Task scans had a rest condition and four contextual conditions: *sur, iso90, orth*, and *iso*. In the rest condition, only a grating was shown inside the fixation circle, excluding target or surround gratings. In the *iso* condition, targets shared orientation and phase with the surround grating. In the *orth* condition, targets had the same phase but orthogonal orientation to the surround. In the *iso90* condition, the targets and surround shared the same orientation but had a relative phase shift of 90°. In the surround-only condition, target gratings were absent, replaced with gray circles matching background luminance (Fig. 1A). All 12-second condition blocks, rest included, were shown eight times in pseudorandom order in a task scan, at each of eight average orientations. Each task scan lasted 480 seconds, and there were four task scans per session.

Localizer scans identified regions of interest for the target and surround. They featured two conditions: target-only (*tgt*) and surround-only (*sur*). In the *tgt* and *sur* conditions, the surround grating or target gratings were absent respectively. As in the task scans, participants performed the 2AFC orientation discrimination task on the center grating (Fig. 1A) and gratings were presented at each of eight orientations. Each 204-second scan began and concluded with a surround block, cycling between surround and target for a total of nine surround and eight target blocks (Fig. 1A). Due to the additional surround block, one orientation appeared in two surround blocks. All other timings and stimuli specifications were consistent between task and localizer scans. In all sessions, three localizer scans were conducted.

### Data collection

Functional MRI data were collected at the University of Minnesota’s Center for Magnetic Resonance Research using a Siemens 7T scanner equipped with a custom-made head coil with a 32-channel transmit and 4-channel receive (Adriany et al., 2012). We employed T_2_*-weighted gradient echo (GE) echo-planar imaging (EPI) (repetition time (TR): 2 s, echo time (TE): 32.2 ms) over 25 coronal slices with a field of view covering the occipital lobe’s posterior extent. The images boasted a 0.6 mm isotropic resolution with a field of view of 124.8 mm x 153.6 mm and a matrix size of 208 x 256. We used an in-plane parallel imaging acceleration factor (R) of 3 with GRAPPA (Griswold et al., 2002) with a right-left phase-encode direction (6/8 Partial Fourier, echo-spacing: 1.2 ms).

During each session, we acquired a T_1_-weighted GE EPI (T_1_wEPI) scan to delineate gray matter in the functional data (van der Zwaag et al., 2018). A reverse phase encode T_1_wEPI scan was used for distortion compensation. Due to the identical resolution, sampling, and echo spacing of the T_1_wEPI and functional data, they were subjected to matching distortions.

Outside this primary session, a T_1_-weighted MP-RAGE scan with 0.8-mm isotropic resolution was obtained on a Siemens 3T scanner. Additionally, seven of the sixteen participants participated in a separate retinotopic mapping scan, either conducted at 3T (two participants) or at 7T (five participants). Retinotopy was acquired using a combination of oriented drifting bars (Dumoulin & Wandell, 2008) and checkerboard-patterned wedges or rings (Engel, Glover, & Wandell, 1997).

Voxel responses to these stimuli were used to fit a population receptive field (pRF) model to each voxel. The resulting polar angle maps were used to refine early visual area boundaries (V1, V2, and V3) in these participants.

### Data pre-processing

#### 3T data

We utilized FreeSurfer’s *recon-all* command to segment the 3T reference anatomy, delineating the GM/white matter (WM) boundary and the pial surface. We used the *hires* expert option to enable adequate surface inflation for submillimeter data, which is necessary for correct surface mapping onto a sphere and subsequent error correction. The cortical surface was then inflated for a maximum of 50 iterations (https://surfer.nmr.mgh.harvard.edu/, v6.00.0).

#### NOise Reduction with DIstribution Corrected (NORDIC) PCA

NORDIC is an algorithm for reducing thermal noise in functional (Vizioli et al., 2021) and diffusion-weighted (Moeller et al., 2021) MRI, which at high resolution is the dominant source of noise. We applied this algorithm to our raw functional data, prior to any other preprocessing, to improve signal-to-noise ratio (SNR), which increased by a factor of 2.14 *±* 0.20. Intrinsic image blurring was estimated before and after processing with NORDIC. Full-width half-maximum (FWHM) of the spatial autocorrelation function of the brain-masked data was estimated from the first localizer scan for each subject before and after applying NORDIC using AFNI’s *3dFWHMx* function. The average effective FWHM of the spatial autocorrelation function across participants was 0.58 *±* 0.02 mm before and 0.58 *±* 0.04 mm after, indicating that no significant spatial correlations were introduced to the data during denoising.

### Functional data

We utilized AFNI tools (https://afni.nimh.nih.gov/afni, v22.0.10) for functional data processing. The *3dvolreg* command facilitated motion compensation, aligning all functional data with the forward phase T_1_wEPI. This procedure was done in two stages: 1) motion compensation within individual scans and 2) motion compensation between scans.

Voxel displacements for distortion compensation were determined with *3dQwarp*, using both forward and reverse T_1_wEPIs. This WARP volume was then merged with motion correction parameters, resulting in a singular re-sampling matrix. This matrix facilitated the simultaneous application of motion and distortion corrections to the functional data, minimizing image blurring from interpolation.

Prior to distortion compensation of the two T_1_wEPI scans and their subsequent averaging, we processed the T_1_wEPI data by fitting the intensity of each voxel based on the slice-specific inversion time for each volume capture. These fits yielded an image with T_1_-weighted contrast, which served as our target for aligning 3T anatomy using *3dAllineate*.

### Statistical Analyses

#### General linear model (GLM) analysis

We employed AFNI’s *3dDeconvolve* for data analysis using a general linear model (GLM). Stimulus regressors were generated by convolving boxcar functions with a standard hemodynamic response function (*HRF* = *t*^4^*e*^−*t*^*/*(4^2^*e*^−4^)), with boxcars aligned to stimulus block onsets and offsets. The design matrix incorporated nuisance regressors (Legendre polynomials up to the fourth-order) to account for baseline drift and motion (from AFNI’s *3dvolreg*). Using the ordinary least squares method, we estimated beta weights. Two distinct GLMs were used, one for localizer scans and the other for task scans. For localizer scans, the design matrix contained regressors for target-only (*tgt*) and surround-only (*sur*) conditions. The task scan GLM estimated response amplitude for the four contextual conditions: *iso, iso90, orth*, and *sur*. Voxels not significantly influenced by the visual stimulus presentation were omitted from subsequent analyses (stimulus vs. rest contrast *p <* 0.01 without correction for multiple comparisons). The average fraction of visually non-responsive voxels excluded from each participant was 18.99 *±* 5.96% with no participant exceeding a voxel dropout rate of 31%.

#### Target ROI delineation

For participants with available retinotopic data, the polar angle maps served as a tool to verify and manually refine the boundaries of V1, V2, and V3, as determined by a publicly available brain atlas (Benson et al., 2014). Atlas demarcations for V1, V2, and V3 were used for all remaining participants who did not have available retinotopic maps. AFNI’s *3dAllineate* was used to align and resample these retinotopically defined V1/V2/V3 ROIs to the in-session T_1_wEPI from 7T. Because the retinotopic data were acquired in a separate scanning session at a lower resolution, we did not directly use the pRF estimates in our localization of voxels to the target region. Instead, the functional localizer scan, which included *tgt* and *sur* conditions, helped in pinpointing the voxels responsive to either the target or the surround. This was achieved by implementing a GLM (outlined above) with dedicated regressors for both conditions.

Target-selective ROIs were created by first defining a cortical surface patch on the FreeSurfer-derived cortical surface mesh centered on the target-selective regions. To do this, the *tgt* - *sur* contrast was projected onto the surface mesh, from which we visually estimated a center point, using the V1 boundary as a reference. A custom Python script then selected the surface nodes lying within a 10mm radius of this point, thereby creating a surface patch that is flattened by FreeSurfer’s *mris flatten* algorithm. This patch then underwent visual inspection to ascertain that all target-selective voxels were incorporated within the patch. The flat patches were then projected back to volume space for analysis. Cortical ribbon coordinates were determined separately in volume space using LayNii’s *LN2_MULTILATERATE* (L. R. Huber et al., 2021) *which were used for ROI definition and visualization (see section on Depth-Dependent Analysis). Unlike standard surface projections which can introduce distoritions to voxel locations, LN2_MULTILATERATE* preserves topographical qualities by operating entirely in volume space. Coordinates, *u* and *v* are assigned to voxels by tracing a path through the middle layer of the cortical ribbon. Together with, the depth coordinate, this creates a flattened cylindrical coordinate system for the cortical ribbon.

Subsequently, the coordinates indicating voxel position along the cortical ribbon were utilized to delineate an elliptical boundary around target-selective voxels within the 10mm patch. First, a 2mm full-width half-max Gaussian blurring kernel was applied to the localizer data to smooth out local variations in the BOLD responses using AFNI’s *3dmerge* command. Next, to identify a region dominated by target-selective responses, we applied a threshold based on the GLM estimate of the target-surround contrast (*p <* 0.01 without correction for multiple comparisons). This leaves a subset of strongly target-selective voxels in the 10 mm patch. Voxels are then projected onto a 2D coordinate space that follows the contours of the middle layer of the cortical ribbon (see section on Depth Dependent Analysis) and the principal components of the spatial covariance matrix are used as the major and minor axes of an elliptical boundary in the flattened cortical space, as shown in Figure 1C. Radial distance is expressed in units of *σ*, which is defined as one standard deviation of the spatial voxel distribution along each principal component. This is found by the square root of the eigenvalues of the spatial covariance matrix. This process offered dual benefits: a refined localization of the demarcation between voxels selective for the target and those for the surround, and a metric of radial distance from the center of the target-selective area to its boundary for each voxel.

We noted that voxels gradually transition from target-selective to surround-selective between 1*σ* and 3*σ*. Therefore, we expected that many voxels in this range likely had pRFs that overlapped with the target edge. To restrict our analysis to voxels that had pRFs well centered on the target, we calculated a theoretical maximum radial distance in voxel pRFs would nearly universally be contained within the target. First, we estimated that the maximum pRF size of voxels representing visual space at 3.75° eccentricity is approximately 1.5° FWHM based on results from Poltoratski & Tong (Poltoratski & Tong, 2020). Due to the lower spatial resolution used by Poltoratski & Tong (Poltoratski & Tong, 2020), our estimate likely slightly overestimates the pRF size of voxels in our analysis. Based on this estimated maximum, pRFs localized to within 1° of the 2.5° target stimulus should not significantly overlap with the edge of the target. We then used the standard formula for cortical magnification in V1 (Horton & Hoyt, 1991), to estimate the scale of the cortical representation of the 1° region:

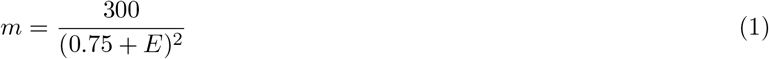

where *m* is the cortical magnification in mm^2^/deg^2^ and *E* is eccentricity. Thus, the cortical magnification at 3.75° eccentricity is approximately 14.81 mm^2^/deg^2^. Restricting our analysis to the region of cortex representing the central 1° of the target grating, this yielded an expected cortical representation size of 11.64 mm^2^. Setting the ROI boundary at 1*σ* resulted in a close match with the expected size from cortical magnification with an average area of 11.93 *±* 4.97 mm^2^. Therefore, we chose to delineate our target-selective ROIs at 1*σ*.

### Depth-dependent analysis

Voxels were given a normalized depth value ranging from 0 to 1, grounded in an equivolume solution within surface space as delineated by Waehnert et al. (Waehnert et al., 2014). This assignment was facilitated using LayNii, a toolset that can be accessed via its GIT repository (https://github.com/layerfMRI/LAYNII) and is documented by Huber et al. (L. R. Huber et al., 2021).

Within each extracted volume patch (see Target ROI delineation), gray matter (GM) and white matter (WM) surface boundaries were drawn manually. The in-session T_1_w EPI was upsampled to 0.3 mm isotropic resolution and overlaid with the FreeSurfer-derived GM and WM segmentations, which were used as guides for manual segmentation. LayNii’s *LN2_RIMIFY* was used to convert the resulting manual segmentation files to rim files, demarcating the GM and WM boundaries in a LayNii-legible format. *LN2_LAYERS* then designated a normalized depth value to each voxel within the ROI. Then *LN2_MULTILATERATE* was used to map coordinates of voxels to their locations along the cortical ribbon. Control points for *LN2_MULTILATERATE* were chosen from the middle GM files produced by *LN2_LAYERS*. The center control point was chosen to be the center of mass of the ROI, and the axes were chosen programmatically to align to orthogonal directions in the FreeSurfer surface coordinate space. However, due to mismatches between FreeSurfer and LayNii’s estimations of the cortical surface, we adjusted the control points manually as needed to ensure that axes were orthogonal and included the full ROI. Finally, the layer and multilaterate files were resampled back to the original functional resolution.

To obtain depth-dependent BOLD profiles, voxels were assigned to seven depth bins of equal width. Voxel responses were averaged within each depth bin and within each participant to obtain depth profiles. Our choice of seven bins was based on the observation that the distribution of voxel centroids essentially constituted a continuous distribution of depths. Therefore, we chose to bin at the highest resolution that would still provide reliable statistics including at least 5 voxels per depth bin per participant. Note however that these voxel counts are for individual participants. All of our statistics are computed across participants in order to examine common trends across participant depth profiles.

All subsequent statistical analyses were performed using custom Python code. Due to the relatively small sample size and in order to account for the possibility of non-normally distributed data, we report permutation tests. However, we found that for all statistical tests, the permutation test was in agreement with the corresponding parametric statistic given a significance level of *α* = 0.05.

For our contrasts of interest, BIM and OTSS,, we performed one-way ANOVAs and Kruskal-Wallis tests to determine if there was an effect of depth on the BOLD contrast (see Tables S1-S4). We also performed a one-way ANOVA to test for contrast-depth interactions between the BIM and OTSS profiles (see Table S5).

As our main analysis, we ran single-sample permutation tests for all of our condition contrasts to test for depths that were significantly modulated (see Tables S6-S9). Permutations were generated by randomly changing the signs of the individual participant averages and computing the mean across participants for each permutation (4096 permutations per comparison). All *p*-values reported are corrected for multiple comparisons using a Benjamini-Hochberg method (Benjamini & Hochberg, 1995) over all one- and two-sample permutation tests for depth and radial profiles. This method for multiple comparisons accounts for measurements that are not statistically independent. This is true of our depth analyses, because voxel centroids formed a continuous distribution across the cortical depth. This means that any parcellation of cortical depth will necessarily involve some signal sharing between depth bins.

#### Deveining

Gradient echo BOLD acquisitions are sensitive to signal contamination from large veins. Voxels located in superficial regions of gray matter are particularly susceptible to this kind of signal contamination due to their proximity to large pial veins. Veinous voxels colocalize with voxels that have high mean-normalized-variance (MNV) in the residual time series data (Olman, Inati, & Heeger, 2007). When inspecting the data, we discovered that the distribution of MNV resembled a log-normal distribution with a long tail. We also noticed that the most superficial 30% of voxels, while having significantly overlapping MNV distributions with the deepest 10% of voxels for all ROIs, had longer MNV tails than in deep regions (Fig. S4). We suspected that this long tail arose from veinous voxels with larger MNV. Therefore, to reduce the contribution from veinous voxels in average depth profiles, we used a threshold based on the MNV to mask out high-MNV voxels. We used the distribution of the deepest 10% of voxels in each ROI to model the MNV distribution of non-veinous voxels, since they were the farthest from large pial veins and therefore were likely to be the least vein-contaminated. Since the deep voxels were well-approximated by a log-normal distribution, we defined the MNV threshold to be two standard deviations above the mean of the deep voxel log(MNV). In most ROIs, this excluded most of the long tails in the most superficial 30% of voxels (Fig. S4). Deveining resulted in a total dropout rate of 11.03 *±* 6.05% which was heavily skewed toward superficial layers (superficial 23.01 *±* 12.95%, middle 7.58 *±* 5.92%, deep 3.30 *±* 1.85%).

An additional source of systematic noise from layer fMRI is the tendency for radially oriented draining veins to carry signal from deep to superficial layers. Signal measured in gray matter voxels therefore represent a mixture of both local neural activity and neural activity from deeper regions (Markuerkiaga et al., 2016). To compensate for this, we followed a deconvolution method developed by Markuerkiaga et al. (Markuerkiaga, Marques, Gallagher, & Norris, 2021) which uses the approximate point spread function (PSF) across cortical depth to deconvolve beta weights. We used a peak-to-tail ratio (p2t) of 6.3 for the PSF which best fit data from Markuerkiaga et al. (Markuerkiaga et al., 2021) acquired at 7T with TE = 33.3 ms, which most closely matched our acquisition. As expected, deconvolution significantly reduced the measurements of average signal in superficial layers relative to deep layers.

#### Radial profiles

To examine the localization of context modulation on the cortical surface, we visualized BOLD contrast as a function of radial distance from the center of the ROI. BOLD responses across the radial axis were convolved with an exponential kernel 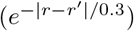 to generate smoothed radial profiles for visualization. We also computed average BOLD contrasts within five discrete radial bins ranging from 0 to 5*σ* at each of three depths.

Single-sample permutation tests were performed for each of three context modulation contrasts (BIM, OTSS, and *iso-sur*) for each contrast within each radial bin (see Tables S13-S15). As with the depth-dependent analysis, a null distribution was obtained by randomly reassigning the signs of the individual participant averages and computing the mean across participants (4096 permutations per comparison). To determine whether voxels localized to the target stimulus were significantly more modulated than voxels well outside the target region, we also statistically compared the center radial bin (0*σ* - 1*σ*) to the most distal radial bin (4*σ* - 5*σ*) for both context modulation contrasts using a two-sample paired permutation test (see Tables S16 and S17). The null distribution was determined by randomly changing the signs of the paired differences within participants and computing the mean across participants. As with previous analyses we used a Benjamini-Hochberg correction for multiple comparisons (Benjamini & Hochberg, 1995) across all one- and two-sample permutation tests for depth and radial profiles.

#### Inclusion criteria

Inclusion criteria were applied to each ROI separately. Identification of ROIs required the presence of a visible target-selective patch in peripheral V1 as determined by the independent functional localizer. We visually assessed both the *tgt* - *sur* contrast on the cortical surface and the elliptical fits to target-selective patches to determine whether an ROI could be drawn in each hemisphere. Hemispheres that did not contain a visually identifiable target-selective region or for which the absence of a clear target-surround delineation precluded an adequate elliptical boundary fit were excluded from final analysis. Of the sixteen participants, fourteen participants had visually identifiable target-selective regions in at least one hemisphere. As a further qualification for analysis, V1 ROIs were required to be significantly target-selective with a minimum of 50% of voxels attaining a significance level of *p <* 0.01 (single-voxel, uncorrected) on the independent functional localizer *tgt-sur* contrast. This criterion resulted in the exclusion of an additional two participants. As a result of these inclusion criteria, we analyzed twenty-one ROIs in twelve out of sixteen participants (see Table S29). All ROIs included in the final analysis had a mean temporal SNR (tSNR) above 10 after distortion compensation. For participants that retained ROIs in both hemispheres, we report results for the combined right and left hemisphere ROIs, yielding *N* = 12 for all analyses. Only significantly stimulus-driven voxels were considered in subsequent analyses. This was determined by assessing the statistical significance of the combined stimulus vs. rest contrast from the task GLM, using a significance threshold of *p <* 0.01. For depth-dependent analysis, we ensured that all included ROIs retained at least 5 voxels in each depth bin.

## Data and Code Availability

Custom code for analyses is written in Python. Omnibus tests (ANOVA and Kruskal-Wallis tests) were performed in JASP https://jasp-stats.org/. Upon publication, data will be made available on OpenNeuro (https://openneuro.org/), and all analysis code will be made available in a public GitHub repository (https://github.com/emers245/OriSeg_analysis).

