## Supplementary Materials for "Orientation-tuned surround suppression exhibits a unique laminar signature in human primary visual cortex"

APRIL 30, 2026

#### Effect of Deconvolution on Laminar Profiles

In the cortical gray matter, draining veins and venules carry deoxygenated blood from the white matter toward the pial surface. This causes signal contamination between cortical depths, leading to a superficial bias in the BOLD signal, whereby superficial layers pool signal from deeper layers leading to stronger observed BOLD contrast (Markuerkiaga et al., 2016). To compensate for this signal leakage, we use a method devised by Markuerkiaga, Marques, Gallagher, and Norris (2021). In their method, Markuerkiaga et al. (2021) estimate the point-spread function (PSF) of the BOLD signal across cortical depth. Given this PSF,

Markuerkiaga et al. (2021) model the BOLD signal observed at a specific depth as a combination of the activation at that depth, plus some fraction of the activation at lower depths. They find that for discretized cortical depth, the depth-dependent PSF is well-approximated by two values that characterize a peak, at the site of activation, and a tail of constant value projecting from the cite of activation toward the pial surface. This is achieved by designing a general linear model (GLM) using the depth-dependent PSFs as regressors to estimate the uncontaminated BOLD depth profile from the original data.

As described in the methods, we use a peak-to-tail ratio (*p2t*) of 6.3 for our depth-dependent regressors. This was found to be the optimal *p2t* for an echo time of TE = 33.3 ms, which was closest to our echo time of TE = 32.2 ms. In our depth deconvolution GLM, the beta weights estimated the uncontaminated BOLD signal at each depth bin given our beta weights estimated from our task and localizer GLMs:

$$\mathbf{B_{experiment}} = \mathbf{X}\mathbf{B_{deconvolved}} + \mathbf{E} \tag{1}$$

where $\mathbf{B_{experiment}}$ is a 7x1 vector containing a single participant's average BOLD percent change for a given condition, $\mathbf{X}$ is a 7x7 design matrix, $\mathbf{B_{deconvolved}}$ is a 7x1 vector estimating the deconvolved signal, and $\mathbf{E}$ is a 7x1 vector of the residual error. For simplicity, we did not account for partial volume effects at the pial surface which would have introduced an additional free parameter into the depth deconvolution GLM.

We compared profiles before and after deconvolution. We noted a clear superficial bias in the profiles prior to deconvolution in the task conditions (Fig. S2). After deconvolution, this superficial bias was significantly reduced (Fig. S2).

Since contrasts between conditions evaluate depth-specific differences in signal, they are less susceptible to the superficial bias observed in condition-versus baseline profiles. However, we still observed a superficial bias in the contrasts prior to deconvolution, but when evaluating contrast based on the deconvolved profiles, the superficial bias was reduced (Fig. S3). We note that all depths, except for the deepest bin remain significantly modulated by the *tgt* stimulus above the *sur* stimulus both before and after depth deconvolution (Fig. S3A,G). We also note that there are no changes in the statistical significance of the *iso-sur* profile before and after deconvolution (Fig. S3D,J). However, there are some changes in the BIM (Fig. S3B,H) and OTSS (Fig. S3C,I) profiles depending on whether depth deconvolution is applied. Specifically, the BIM contrast for the bin centered at 0.67 and the OTSS contrast for the bin centered at 1.00 are statistically significant prior to deconvolution. However, these shifts caused by depth deconvolution do not significantly impact our interpretation of the results. Even before deconvolution, we find local peak BIM contrast in deep and super and superficial bins and peak OTSS contrast in superficial bins.

For thoroughness, we analyzed two additional contrasts: *orth–iso* and *tgt–iso*. The *orth–iso* contrast reflects the combined influence of BIM and OTSS. Depth deconvolution did not change any depth-dependent statistical outcomes for this contrast (Fig. S3E,K). However, because it combines multiple mechanisms, we found it more informative to analyze the BIM and OTSS profiles separately. We also explored a *tgt–iso* contrast to estimate the overall magnitude of surround suppression. Interpreting this contrast proved more challenging: the *tgt* and *iso* responses were obtained in different experimental runs and estimated using separate GLMs. Whereas task conditions were modeled relative to a blank baseline, the localizer GLM contrasted only the two stimulus conditions without a blank. To place the *iso* estimates within the same reference frame as the localizer (*tgt* and *sur*), we subtracted the task GLM *sur* beta weights from the *iso* beta weights and then contrasted these adjusted values with the localizer GLM *tgt* beta weights. Depth deconvolution substantially altered the resulting *tgt–iso* profile, with significant effects appearing only in deep bins after deconvolution. Moreover, these profiles were markedly more variable, especially in superficial layers, than contrasts derived from conditions estimated within the same GLM (Fig. S3F,L), despite having similar average magnitudes to the *tgt–sur* contrast. We attribute this variability to the differing baselines in the task and localizer scans. Consequently, we do not consider the *tgt–iso* profiles to provide reliable indicators of surround suppression across depth.

### Deveining

Vein masks were constructed using the mean-normalized variance (MNV) of the residual time series data. Since veins colocalize with high MNV (Olman et al., 2007), voxels exceeding a

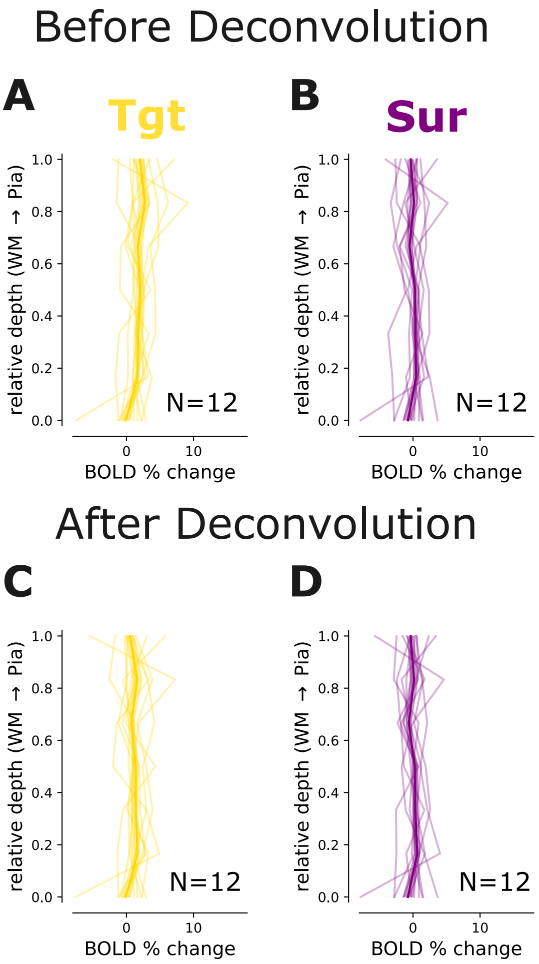


Supplemental Figure 1: **Localizer Depth Deconvolution.** (A-B) Depth profiles for *tgt* (A) and *sur* (B) conditions. Light-colored lines indicate individual subject averages within the target-selective ROIs. Bold line indicates grand average over all subjects. Shaded region indicates standard error of the mean. All profiles are shown prior to deconvolution. (C-D) Following the same conventions as A-B, the depth profiles for both *tgt* (C) and *sur* (D) conditions are shown after depth deconvolution.

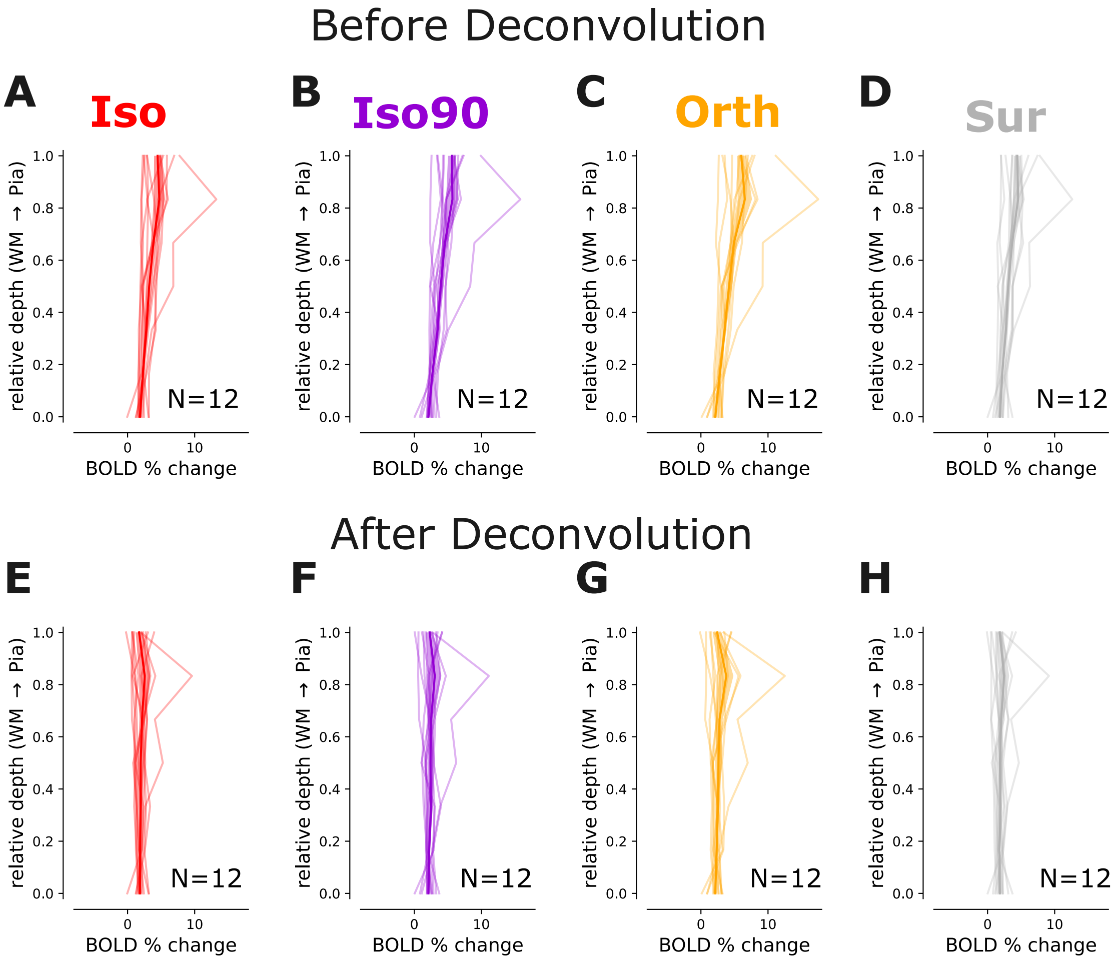


Supplemental Figure 2: **Task Depth Deconvolution.** (A-D) Depth profiles for *iso* (A), *iso90* (B), *orth*, and *sur* task conditions. Light-colored lines indicate individual subject averages within the target-selective ROIs. Bold line indicates grand average over all subjects. Shaded region indicates standard error of the mean. All profiles are shown prior to deconvolution. (E-H) Following the same conventions as A-D, the depth profiles for *iso* (E), *iso90* (F), *orth* (G), and *sur* (H) conditions are shown after depth deconvolution.

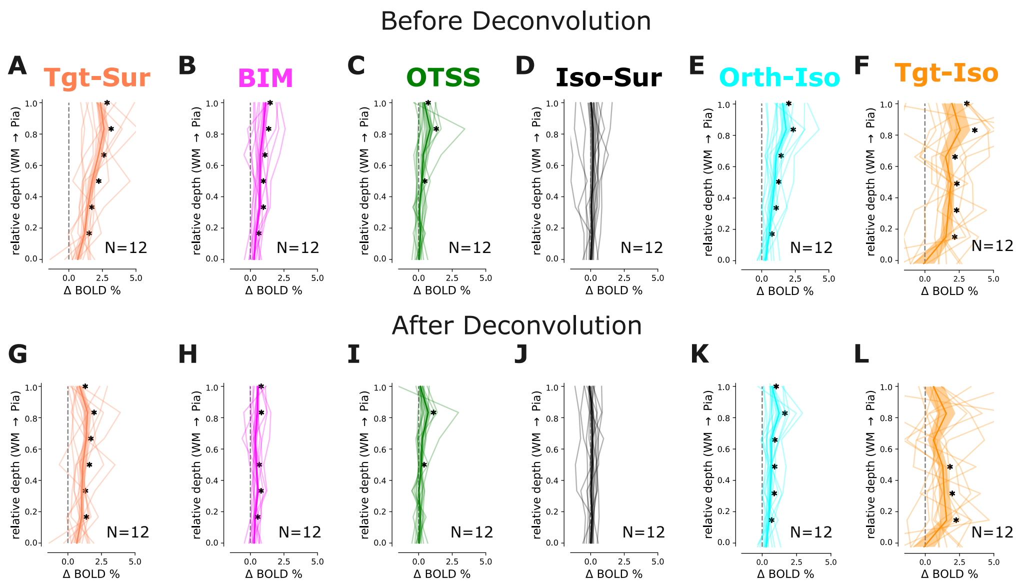


Supplemental Figure 3: **Effects of Depth Deconvolution on Condition Contrasts.** (A-F) Depth profiles for *ctr-sur* (A), BIM (*iso90-iso*) (B), OTSS (*orth-iso90*) (C), *iso-sur* (D), *orth-iso* (E), and *tgt-iso* (F) condition contrasts. Light-colored lines indicate individual subject averages within the target-selective ROIs. Bold line indicates grand average over all subjects. Shaded region indicates standard error of the mean. All profiles are shown prior to deconvolution. (G-L) Following the same conventions as A-F, the depth profiles for *ctr-sur* (G), BIM (*iso90-iso*) (H), OTSS (*orth-iso90*) (I), *iso-sur* (J), *orth-iso* (K), and *tgt-iso* (L) condition contrasts are shown after depth deconvolution. See Tables S6 - S11 for statistics.

mean-normalized variance threshold, specified independently for each ROI, were excluded from analysis (see Methods for details). The MNV distributions for each ROI are shown in Figure S4. While the median log(MNV) was similar between deep and superficial bins, superficial bins were more likely to show long tails of high log(MNV) voxels (Fig. S4A & D). These voxels with high MNV were also highly colocalized suggesting a common veinous origin (Fig. S4B). As described in the methods, setting a threshold at the 95th percentile of the deep bin log(MNV) distribution excluded a greater fraction of voxels in superficial bins where the distributions were more likely to reflect a combination of veinous and non-veinous signals.

### Depth Statistics

Depth profiles were created for five condition contrasts: localizer (*tgt-sur*), BIM (*iso90-iso*), OTSS (*orth-iso90*), *iso-sur*, and *orth-iso*. To tests for effects of depth on condition contrasts BIM and OTSS, we used both one-way ANOVAs (Tables S1 & S3) and Kruskal-Wallis tests (Tables S2 & S4). We also directly compared BIM and OTSS depth profiles to test for condition contrast-depth interactions using a two-way repeated measures ANOVA (Table S5).

Single-sample two-sided permutation tests were used to assess whether the condition contrast at each depth bin was significantly different than zero. These tests were used to identify which depth bins were significantly modulated by the condition contrasts. All $p$-values were corrected for multiple comparisons using an FDR Benjamini-Hochberg correction (Benjamini & Hochberg, 1995) across all one- and two-sample permutation tests for depth and radial profiles. Tables S6-S10 report statistics for all single-sample permutation tests across depth. See Methods for further details on depth analyses.

Supplemental Table 1: **BIM ANOVA.** One-way ANOVA for BIM depth profile ($N = 12$; Fig. 4D).

| Cases | Sum of Squares | df | Mean Square | F | p |
|---|---|---|---|---|---|
| Depth | 0.967 | 6 | 0.161 | 0.922 | $4.84 \times 10^{-1}$ |

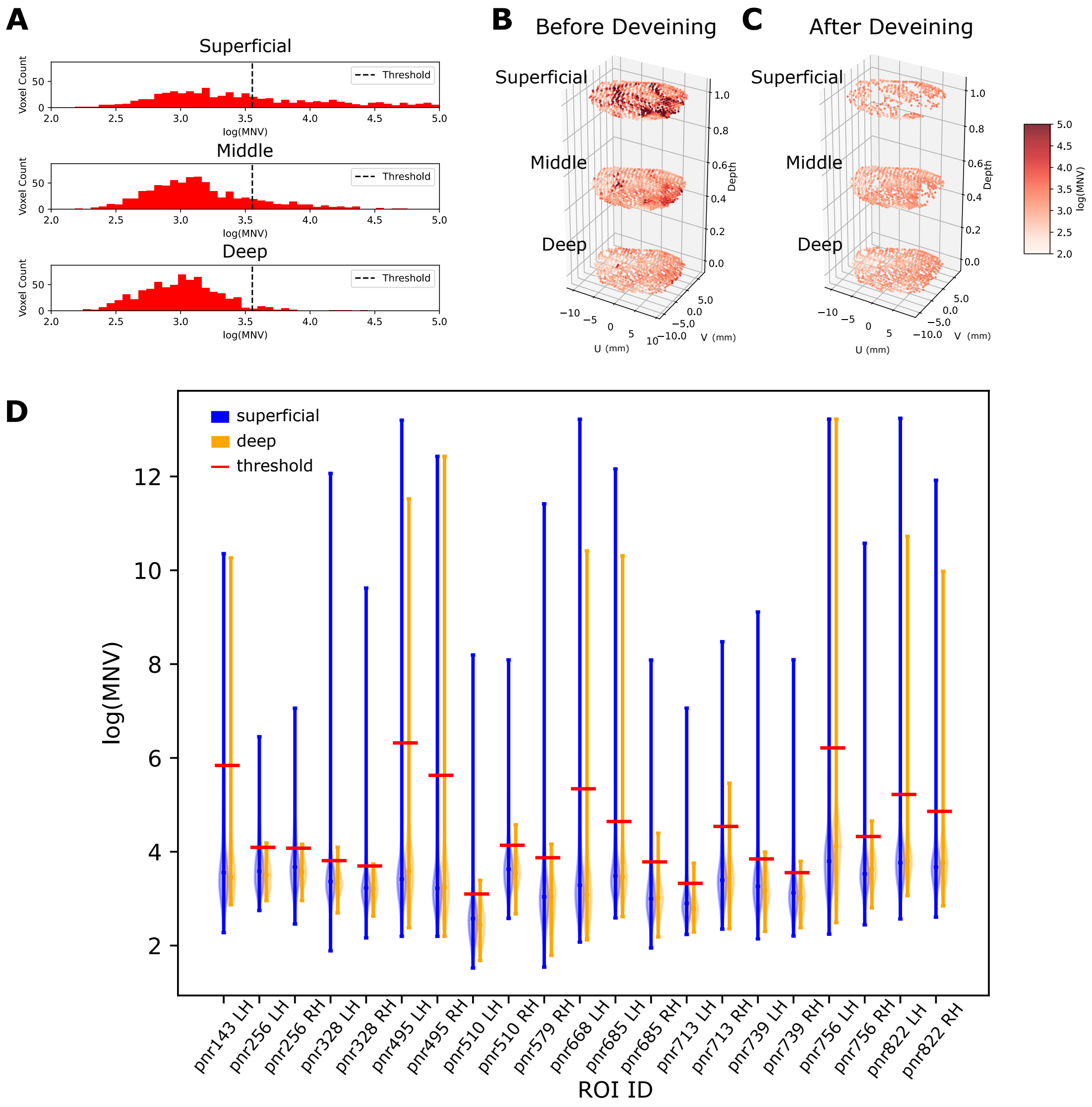


Supplemental Figure 4: **Removal of veinous voxels.** Veinous voxels were identified by analyzing the mean-normalized variance of the residual timeseries for each voxel after the GLM analysis removed stimulus-driven responses. (A) Histograms of the natural logarithm of the mean normalized variance (log(MNV)) of the residual time series are plotted to show the distribution over voxels in an example ROI (pnr739 RH). Histograms are plotted separately for three equally sized depth bins. The dotted line indicates the threshold used for all bins determined from the log(MNV) of the deepest 10% of voxels. (B-C) 3D visualizations of flattened cortical space demonstrates the spatial distribution of log(MNV) at three separate depth bins. Color indicates the log(MNV) for each voxel. (B). After thresholding, voxels with high log(MNV) are removed (C). (D) Violin plots directly compare the distribution of log(MNV) in deep (orange) and superficial (blue) regions. Shaded regions show the probability density of the data, a central horizontal bar indicates the median value, and the tails show the range of the data.

Supplemental Table 2: **BIM Kruskal-Wallis test.** Kruskal-Wallis test for BIM depth profile ($N = 12$; Fig. 4D).

| Cases | statistic | df | p |
|---|---|---|---|
| Depth | 5.15 | 6 | $5.25 \times 10^{-1}$ |

Supplemental Table 3: **OTSS ANOVA.** One-way ANOVA for OTSS depth profile ($N = 12$; Fig. 4E).

| Cases | Sum of Squares | df | Mean Square | F | p |
|---|---|---|---|---|---|
| Depth | 4.11 | 6 | 0.686 | 2.99 | $1.12 \times 10^{-2}$* |

Supplemental Table 4: **OTSS Kruskal-Wallis test.** Kruskal-Wallis test for OTSS depth profile ($N = 12$; ; Fig. 4E).

| Cases | statistic | df | p |
|---|---|---|---|
| Depth | 12.8 | 6 | $4.64 \times 10^{-2}$* |

Supplemental Table 5: **OTSS vs BIM ANOVA.** Two-way repeated measures ANOVA for OTSS vs BIM depth profile ($N = 12$; Fig. 4).

| Cases | Sphericity Corr. | Sum of Sq. | df | Mean Sq. | F | p |
|---|---|---|---|---|---|---|
| Contrast | None | 2.29 | 1.00 | 2.29 | 6.54 | $2.67 \times 10^{-2}$* |
| Depth | None | 2.99 | 6.00 | 0.498 | 5.75 | $7.41 \times 10^{-5}$* |
| | Greenhouse-Geisser | 2.99 | 3.18 | 0.938 | 5.75 | $2.24 \times 10^{-3}$* |
| | Huynh-Feldt | 2.99 | 4.63 | 0.645 | 5.75 | $3.83 \times 10^{-4}$* |
| Contrast * Depth | None | 2.10 | 6.00 | 0.395 | 1.53 | $1.83 \times 10^{-1}$ |
| | Greenhouse-Geisser | 2.10 | 1.80 | 1.17 | 1.53 | $2.42 \times 10^{-1}$ |
| | Huynh-Feldt | 2.10 | 2.12 | 0.97 | 1.53 | $2.38 \times 10^{-1}$ |

Supplemental Table 6: ***tgt-sur* Depth Statistics.** Statistics from single-sample two-sided permutation tests of the *tgt-sur* depth profiles are shown. Averages are taken across all subject profiles ($N = 12$; Fig. 2C & S3F).

| Depth Bin | Normalized Depth | *tgt-sur* Mean ($\Delta$ % BOLD Change) | St. Dev. | *p*-value |
|---|---|---|---|---|
| 0 (deep) | 0.000 | 0.663 | 0.895 | $5.48 \times 10^{-2}$ |
| 1 | 0.167 | 1.02 | 0.680 | $3.73 \times 10^{-3}$* |
| 2 | 0.333 | 1.04 | 0.461 | $2.36 \times 10^{-3}$* |
| 3 | 0.500 | 1.13 | 1.02 | $5.80 \times 10^{-3}$* |
| 4 | 0.667 | 1.36 | 0.695 | $2.36 \times 10^{-3}$* |
| 5 | 0.833 | 1.47 | 1.04 | $2.36 \times 10^{-3}$* |
| 6 (superficial) | 1.00 | 0.884 | 0.856 | $2.36 \times 10^{-3}$* |

Supplemental Table 7: **BIM Depth Statistics.** Statistics from single-sample two-sided permutation tests of the BIM (*iso90-iso*) depth profiles are shown. Averages are taken across all subject profiles ($N = 12$; Fig. 4D & S3G).

| Depth Bin | Normalized Depth | BIM Mean (Δ % BOLD Change) | St. Dev. | $p$-value |
|---|---|---|---|---|
| 0 (deep) | 0.000 | 0.272 | 0.368 | $5.46 \times 10^{-2}$ |
| 1 | 0.167 | 0.363 | 0.243 | $2.36 \times 10^{-3}$* |
| 2 | 0.333 | 0.559 | 0.374 | $2.36 \times 10^{-3}$* |
| 3 | 0.500 | 0.456 | 0.289 | $2.62 \times 10^{-3}$* |
| 4 | 0.667 | 0.361 | 0.530 | $6.49 \times 10^{-2}$ |
| 5 | 0.833 | 0.514 | 0.559 | $1.49 \times 10^{-2}$* |
| 6 (superficial) | 1.00 | 0.584 | 0.334 | $2.62 \times 10^{-3}$* |

Supplemental Table 8: **OTSS Depth Statistics.** Statistics from single-sample two-sided permutation tests of the OTSS (*orth-iso90*) depth profiles are shown. Averages are taken across all subject profiles ($N = 12$; Fig. 4E & S3H).

| Depth Bin | Normalized Depth | OTSS Mean (Δ % BOLD Change) | St. Dev. | $p$-value |
|---|---|---|---|---|
| 0 (deep) | 0.000 | 0.0219 | 0.296 | $8.42 \times 10^{-1}$ |
| 1 | 0.167 | 0.116 | 0.256 | $2.07 \times 10^{-1}$ |
| 2 | 0.333 | 0.0247 | 0.339 | $8.42 \times 10^{-1}$ |
| 3 | 0.500 | 0.212 | 0.248 | $9.37 \times 10^{-3}$* |
| 4 | 0.667 | 0.265 | 0.464 | $7.90 \times 10^{-2}$ |
| 5 | 0.833 | 0.713 | 0.793 | $2.62 \times 10^{-3}$* |
| 6 (superficial) | 1.00 | 0.122 | 0.546 | $5.88 \times 10^{-1}$ |

Supplemental Table 9: ***iso-sur* Depth Statistics.** Statistics from single-sample two-sided permutation tests of the *iso-sur* depth profiles are shown. Averages are taken across all subject profiles ($N = 12$; Fig. 5C & S3I).

| Depth Bin | Normalized Depth | *iso-sur* Mean (Δ % BOLD Change) | St. Dev. | $p$-value |
|---|---|---|---|---|
| 0 (deep) | 0.000 | 0.00849 | 0.335 | $9.35 \times 10^{-1}$ |
| 1 | 0.167 | 0.0995 | 0.247 | $2.67 \times 10^{-1}$ |
| 2 | 0.333 | 0.0260 | 0.487 | $8.89 \times 10^{-1}$ |
| 3 | 0.500 | 0.120 | 0.450 | $4.99 \times 10^{-1}$ |
| 4 | 0.667 | 0.126 | 0.477 | $4.75 \times 10^{-1}$ |
| 5 | 0.833 | 0.0994 | 0.514 | $6.02 \times 10^{-1}$ |
| 6 (superficial) | 1.00 | -0.0747 | 0.482 | $6.83 \times 10^{-1}$ |

Supplemental Table 10: ***orth-iso* Depth Statistics.** Statistics from single-sample two-sided permutation tests of the *orth-iso* depth profiles are shown. Averages are taken across all subject profiles ($N = 12$; Fig. S3J).

| Depth Bin | Normalized Depth | *orth-iso* Mean (Δ % BOLD Change) | St. Dev. | *p*-value |
|---|---|---|---|---|
| 0 (deep) | 0.000 | 0.294 | 0.478 | $8.60 \times 10^{-2}$ |
| 1 | 0.167 | 0.479 | 0.291 | $2.36 \times 10^{-3}$* |
| 2 | 0.333 | 0.583 | 0.506 | $3.73 \times 10^{-3}$* |
| 3 | 0.500 | 0.668 | 0.368 | $2.36 \times 10^{-3}$* |
| 4 | 0.667 | 0.625 | 0.523 | $6.53 \times 10^{-3}$* |
| 5 | 0.833 | 1.23 | 0.858 | $2.36 \times 10^{-3}$* |
| 6 (superficial) | 1.00 | 0.705 | 0.562 | $5.80 \times 10^{-3}$* |

Supplemental Table 11: ***tgt-iso* Depth Statistics.** Statistics from single-sample two-sided permutation tests of the *tgt-iso* depth profiles are shown. Averages are taken across all subject profiles ($N = 12$; Fig. S3L).

| Depth Bin | Normalized Depth | *tgt-iso* Mean (Δ % BOLD Change) | St. Dev. | *p*-value |
|---|---|---|---|---|
| 0 (deep) | 0.000 | -0.0917 | 2.71 | $9.35 \times 10^{-1}$ |
| 1 | 0.167 | 1.56 | 1.59 | $1.29 \times 10^{-2}$* |
| 2 | 0.333 | 1.37 | 1.31 | $7.51 \times 10^{-3}$* |
| 3 | 0.500 | 1.28 | 1.13 | $6.54 \times 10^{-3}$* |
| 4 | 0.667 | 0.603 | 1.52 | $2.70 \times 10^{-1}$ |
| 5 | 0.833 | 1.53 | 2.28 | $6.50 \times 10^{-2}$ |
| 6 (superficial) | 1.00 | 0.601 | 2.67 | $5.34 \times 10^{-1}$ |

Supplemental Table 12: **Voxel Count per Depth Bin.** The average number of significantly visually responsive voxels per participant for the target-selective ROI after deveining is reported for each depth bin.

| Depth Bin | Normalized Depth | Average Voxel Count ± St. Dev. |
|---|---|---|
| 0 (deep) | 0.000 | 7.5 ± 3.5 |
| 1 | 0.167 | 17.1 ± 8.7 |
| 2 | 0.333 | 22.4 ± 11.0 |
| 3 | 0.500 | 23.6 ± 11.9 |
| 4 | 0.667 | 25.0 ± 12.9 |
| 5 | 0.833 | 21.25 ± 11.9 |
| 6 (superficial) | 1.00 | 18.1 ± 11.4 |

### Radial Statistics

Radial profiles were created for three condition contrasts using five radial bins: BIM (*iso90-iso*), OTSS (*orth-iso90*), and *iso-sur*. Single-sample two-sided permutation tests were used to assess whether the condition contrast at each radial bin was significantly different than zero.

All $p$-values were corrected for multiple comparisons using an FDR Benjamini-Hochberg correction (Benjamini & Hochberg, 1995) across all one- and two-sample permutation tests for depth and radial profiles. Statistics for one-sample permutation tests are reported in Tables S13-S15.

For BIM and OTSS, additional two-sample paired permutation tests were used to compare condition contrasts between the central radial bin (0-1$\sigma$) and the outermost bin (4-5$\sigma$). As for depth profiles, $p$-values were corrected for multiple comparisons using an FDR Benjamini-Hochberg correction (Benjamini & Hochberg, 1995) across all one- and two-sample permutation tests for depth and radial profiles. Statistics for two-sample permutation tests between central and outermost bins are reported in Tables S16 & S17. See Methods for further details on radial analyses.

Supplemental Table 13: **BIM Radial Statistics.** Statistics from single-sample two-sided permutation tests of the BIM (*iso90-iso*) radial profiles are shown. Averages are taken across all subject profiles ($N = 12$; Fig. 3D).

| Radius | BIM Mean ($\Delta$ % BOLD Change) | St. Dev. | $p$-value |
|---|---|---|---|
| $0-1\sigma$ | 0.781 | 0.507 | $2.62 \times 10^{-3}*$ |
| $1-2\sigma$ | 0.793 | 0.377 | $2.36 \times 10^{-3}*$ |
| $2-3\sigma$ | 0.845 | 0.358 | $2.36 \times 10^{-3}*$ |
| $3-4\sigma$ | 0.618 | 0.384 | $2.62 \times 10^{-3}*$ |
| $4-5\sigma$ | 0.429 | 0.417 | $9.37 \times 10^{-3}*$ |

Supplemental Table 14: **OTSS Radial Statistics.** Statistics from single-sample two-sided permutation tests of the OTSS (*orth-iso90*) radial profiles are shown. Averages are taken across all subject profiles ($N = 12$; Fig. 3E).

| Radius | OTSS Mean (Δ % BOLD Change) | St. Dev. | $p$-value |
|---|---|---|---|
| $0-1\sigma$ | 0.360 | 0.391 | $2.49 \times 10^{-2}$* |
| $1-2\sigma$ | 0.260 | 0.268 | $1.88 \times 10^{-2}$* |
| $2-3\sigma$ | 0.253 | 0.374 | $5.65 \times 10^{-2}$ |
| $3-4\sigma$ | 0.218 | 0.377 | $7.90 \times 10^{-2}$ |
| $4-5\sigma$ | 0.181 | 0.352 | $1.65 \times 10^{-1}$ |

Supplemental Table 15: ***iso-sur* Radial Statistics.** Statistics from single-sample two-sided permutation tests of the *iso-sur* radial profiles are shown. Averages are taken across all subject profiles ($N = 12$; Fig. 5D).

| Radius | *iso-sur* Mean (Δ % BOLD Change) | St. Dev. | $p$-value |
|---|---|---|---|
| $0-1\sigma$ | 0.121 | 0.673 | $6.18 \times 10^{-1}$ |
| $1-2\sigma$ | -0.0475 | 0.520 | $8.18 \times 10^{-1}$ |
| $2-3\sigma$ | -0.427 | 0.481 | $1.43 \times 10^{-2}$* |
| $3-4\sigma$ | -0.441 | 0.447 | $1.29 \times 10^{-2}$* |
| $4-5\sigma$ | -0.336 | 0.435 | $4.33 \times 10^{-2}$* |

Supplemental Table 16: **BIM Radial Multiple Comparisons.** Statistics from paired two-sided permutation tests of the comparison between central (0-1$\sigma$) and distal (4-5$\sigma$) radial bins for the BIM (*iso90-iso*) contrast are shown. Averages are taken across all subject profiles ($N = 12$; Fig. 3D). Stars indicate statistical significance ($p < 0.05$).

| Bin 1 Radius | Bin 2 Radius | BIM Mean Bin 1 ± St. Dev. (Δ % BOLD Change) | BIM Mean Bin 2 ± St. Dev. (Δ % BOLD Change) | $p$-value |
|---|---|---|---|---|
| $0-1\sigma$ | $4-5\sigma$ | $0.781 \pm 0.507$ | $0.429 \pm 0.417$ | $1.07 \times 10^{-2}$* |

Supplemental Table 17: **OTSS Radial Multiple Comparisons.** Statistics from paired two-sided permutation tests of the comparison between central (0-1$\sigma$) and distal (4-5$\sigma$) radial bins for the OTSS (*orth-iso90*) contrast are shown. Averages are taken across all subject profiles ($N = 12$; Fig. 3E). Stars indicate statistical significance ($p < 0.05$).

| Bin 1 Radius | Bin 2 Radius | OTSS Mean Bin 1 ± St. Dev. (Δ % BOLD Change) | OTSS Mean Bin 2 ± St. Dev. (Δ % BOLD Change) | $p$-value |
|---|---|---|---|---|
| $0-1\sigma$ | $4-5\sigma$ | $0.360 \pm 0.391$ | $0.181 \pm 0.352$ | $2.12 \times 10^{-2}$* |

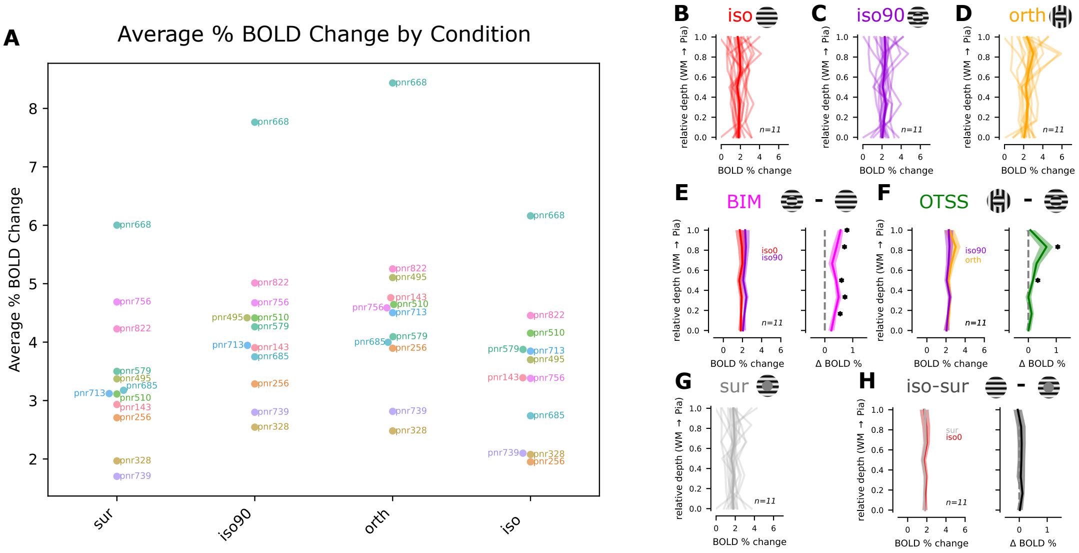


Supplemental Figure 5: **Removal of pnr668 from Depth Analysis:** A) Average percent BOLD change in V1 target ROI ($< 1\sigma$) within each condition. Dots are labeled with participant IDs. B-D) Depth profiles for *iso* (B), *iso90* (C), and *orth* (D) conditions without pnr668. Plots follow the conventions of Fig. 4A-C: within subject averages shown in light-colored lines, and grand averages in bold lines. E-F) Panels show the depth profiles for BIM and OTSS after excluding pnr668. E) Left: Grand average depth profiles for *iso* and *iso90* conditions. Right: Grand average depth profile for the *iso90-iso* contrast (BIM). F) Left: Grand average depth profiles for *orth* and *iso90* conditions. Right: Grand average depth profile for the *orth-iso90* contrast (OTSS). G) Depth profiles for the *sur* condition without pnr668, following the same conventions as panels B-D. F) Left: Grand average depth profiles for *iso* and *sur* conditions. Right: Grand average depth profile for the *iso* - *sur* conditions, following the same conventions as panels E-F. For all panels, shaded regions indicate standard error of the mean across participants. Stars indicate significant release from suppression ($p < 0.05$, single-sample two-sided permutation test, $N_{perm} = 4096$, FDR-corrected, see Tables S23-S25).

### Effects of Outlier on Laminar Profiles

BOLD contrast can differ greatly between individuals. However, we noted one participant, pnr668, with BOLD responses greater than two standard deviations above the mean across participants in all conditions relative to the rest condition (*sur*: $z = 2.25$, *iso90*: $z = 2.65$, *orth*: $z = 2.62$, iso: $z = 2.24$; Fig. S5A). To determine whether this outlier disproportionately influenced the grand average depth profiles, we reran our depth profile analyses after removing this subject, yielding eleven participants. We found that removing this participant did not affect our determination of statistical significance ($\alpha = 0.05$) for our omnibus tests of the BIM profile (Kruskal-Wallis $H = 5.48$, $p = 0.484$; one-way ANOVA $F = 1.09$, $p = 0.378$), showing no significant effect of depth. We also did not observe any changes in our determination of statistical significance for pairwise permutation tests for BIM (*iso90* vs *iso*), OTSS (*orth* vs *iso90*), or *iso-sur* (Fig. S5B-F; Table S23-S25). However, removing pnr668 returned mixed results for the omnibus tests of the OTSS profile, with a significant effect of depth being maintained in the one-way ANOVA ($F = 2.45$, $p = 0.033$) but not the Kruskal-Wallis test (Kruskal-Wallis $H = 11.52$, $p = 0.074$). Furthermore, while the main effect of depth remained in the comparison between BIM and OTSS profiles (two-way repeated measures ANOVA $F = 4.27$, $p = 0.001$), the effect of condition contrast shifted slightly above our significance threshold (two-way repeated measures ANOVA $F = 4.66$, $p = 0.056$). The interaction between depth and condition contrast remained insignificant (two-way repeated measures $F = 1.60$, $p = 0.162$). Despite these slight changes to the significance levels of the omnibus tests, the trends remained consistent and our main conclusions did not change: OTSS is present in superficial but not deep layers, while BIM is broadly distributed across depth. We also did not observe any large qualitative shifts in the depth profiles. Since pnr668 already met all inclusion criteria, we include this participant in our main manuscript.

Supplemental Table 18: **BIM ANOVA without pnr668.** One-way ANOVA for BIM depth profile after removing pnr668 ($N = 11$; Fig. S5E).

| Cases | Sum of Squares | df | Mean Square | F | p |
|---|---|---|---|---|---|
| Depth | 0.934 | 6 | 0.156 | 1.09 | $3.78 \times 10^{-1}$ |

Supplemental Table 19: **BIM Kruskal-Wallis test without pnr668.** Kruskal-Wallis test for BIM depth profile after removing pnr668 ($N = 11$; Fig. S5E).

| Cases | statistic | df | p |
|---|---|---|---|
| Depth | 5.477 | 6 | $4.84 \times 10^{-1}$ |

Supplemental Table 20: **OTSS ANOVA without pnr668.** One-way ANOVA for OTSS depth profile after removing pnr668 ($N = 11$; Fig. S5F).

| Cases | Sum of Squares | df | Mean Square | F | p |
|---|---|---|---|---|---|
| Depth | 3.43 | 6 | 0.572 | 2.45 | $3.31 \times 10^{-2}$* |

Supplemental Table 21: **OTSS Kruskal-Wallis test without pnr668.** Kruskal-Wallis test for OTSS depth profile after removing pnr668 ($N = 11$; ; Fig. S5F.

| Cases | statistic | df | p |
|---|---|---|---|
| Depth | 11.5 | 6 | $7.37 \times 10^{-2}$ |

Supplemental Table 22: **OTSS vs BIM ANOVA without pnr668.** Two-way repeated measures ANOVA for OTSS vs BIM depth profile after removing pnr668 ($N = 11$; Fig. S5).

| Cases | Sphericity Corr. | Sum of Sq. | df | Mean Sq. | F | p |
|---|---|---|---|---|---|---|
| Contrast | None | 1.62 | 1.00 | 1.62 | 4.66 | $5.63 \times 10^{-2}$ |
| Depth | None | 2.10 | 6.00 | 0.349 | 4.27 | $1.21 \times 10^{-3}$* |
| | Greenhouse-Geisser | 2.10 | 2.96 | 0.709 | 4.27 | $1.31 \times 10^{-2}$* |
| | Huynh-Feldt | 2.10 | 4.33 | 0.484 | 4.27 | $4.41 \times 10^{-3}$* |
| Contrast * Depth | None | 2.27 | 6.00 | 0.378 | 1.60 | $1.62 \times 10^{-1}$ |
| | Greenhouse-Geisser | 2.27 | 1.68 | 1.35 | 1.60 | $2.31 \times 10^{-1}$ |
| | Huynh-Feldt | 2.27 | 1.98 | 1.15 | 1.60 | $2.27 \times 10^{-1}$ |

Supplemental Table 23: **BIM Depth Statistics without pnr688.** Statistics from single-sample two-sided permutation tests of the BIM (*iso90-iso*) depth profiles after removing pnr668 are shown. Averages are taken across all remaining subject profiles ($N = 11$; Fig. S5E).

| Depth Bin | Normalized Depth | BIM Mean (Δ % BOLD Change) | St. Dev. | *p*-value |
|---|---|---|---|---|
| 0 (deep) | 0.000 | 0.234 | 0.361 | $1.04 \times 10^{-1}$ |
| 1 | 0.167 | 0.356 | 0.252 | $4.57 \times 10^{-3}$* |
| 2 | 0.333 | 0.497 | 0.328 | $4.57 \times 10^{-3}$* |
| 3 | 0.500 | 0.406 | 0.249 | $5.06 \times 10^{-3}$* |
| 4 | 0.667 | 0.263 | 0.437 | $1.23 \times 10^{-1}$ |
| 5 | 0.833 | 0.422 | 0.487 | $2.90 \times 10^{-2}$* |
| 6 (superficial) | 1.00 | 0.565 | 0.342 | $5.06 \times 10^{-3}$* |

Supplemental Table 24: **OTSS Depth Statistics without pnr668.** Statistics from single-sample two-sided permutation tests of the OTSS (*orth-iso90*) depth profiles after removing pnr668 are shown. Averages are taken across all remaining subject profiles ($N = 11$; Fig. 5F).

| Depth Bin | Normalized Depth | OTSS Mean (Δ % BOLD Change) | St. Dev. | *p*-value |
|---|---|---|---|---|
| 0 (deep) | 0.000 | 0.00446 | 0.303 | $9.79 \times 10^{-1}$ |
| 1 | 0.167 | 0.117 | 0.267 | $2.55 \times 10^{-1}$ |
| 2 | 0.333 | 0.00272 | 0.345 | $9.87 \times 10^{-1}$ |
| 3 | 0.500 | 0.164 | 0.199 | $1.82 \times 10^{-2}$* |
| 4 | 0.667 | 0.293 | 0.475 | $8.22 \times 10^{-2}$ |
| 5 | 0.833 | 0.651 | 0.799 | $5.06 \times 10^{-3}$* |
| 6 (superficial) | 1.00 | 0.0739 | 0.546 | $7.77 \times 10^{-1}$ |

Supplemental Table 25: ***iso-sur* Depth Statistics without pnr668.** Statistics from single-sample two-sided permutation tests of the *iso-sur* depth profiles after removing pnr668 are shown. Averages are taken across all remaining subject profiles ($N = 11$; Fig. 5H.

| Depth Bin | Normalized Depth | *iso-sur* Mean (Δ % BOLD Change) | St. Dev. | *p*-value |
|---|---|---|---|---|
| 0 (deep) | 0.000 | 0.0208 | 0.348 | $8.85 \times 10^{-1}$ |
| 1 | 0.167 | 0.113 | 0.253 | $2.50 \times 10^{-1}$ |
| 2 | 0.333 | 0.0550 | 0.499 | $7.77 \times 10^{-1}$ |
| 3 | 0.500 | 0.0765 | 0.445 | $7.22 \times 10^{-1}$ |
| 4 | 0.667 | 0.0863 | 0.478 | $6.74 \times 10^{-1}$ |
| 5 | 0.833 | 0.0673 | 0.525 | $7.57 \times 10^{-1}$ |
| 6 (superficial) | 1.00 | -0.0559 | 0.499 | $7.77 \times 10^{-1}$ |

#### Orientation Decoding Analysis

To further assess whether the responses in the target ROI during the *sur* condition were due to a neural origin or to blurring from the surrounding cortex, we performed an orientation-decoding analysis using a support vector machine (SVM). We reasoned that spatial spread of the BOLD signal might carry surround orientation information to superficial layers but not deep layers, while neural mechanisms could communicate surround orientation at all depths. We implemented the SVM with scikit-learn's *svm.SVC* function `https://scikit-learn.org/stable/modules/generated/sklearn.svm.SVC.html`. The SVM solves an eight-way classification problem to decode stimulus orientation from voxel responses at TRs 2-6 (4-12 seconds) after stimulus onset. The multiclass SVM is implemented using a one-vs-one strategy, yielding 28 binary classifiers trained to distinguish between each pair of orientations. Each binary classifier can be expressed in the Lagrangian dual form:

$$f(\mathbf{x}) = \sum_{i \in \mathrm{SV}} \alpha_i y_i K(\mathbf{x_i}, \mathbf{x}) + b \tag{2}$$

where $f(\mathbf{x})$ is a decision function evaluated for a voxel response vector $\mathbf{x}$. The classifier's decision boundary is defined by a linear combination of support vectors (training samples lying closest to the boundary), with corresponding weights $\alpha_i$, class labels $y_i \in \{-1, 1\}$, and bias $b$. The SVM maximizes the margin between classes while penalizing misclassified samples according to a regularization parameter $C$, which controls the trade-off between margin width and classification error.

We use a radial basis function (RBF) kernel to permit nonlinear decision boundaries:

$$K(\mathbf{x}_i, \mathbf{x}) = \exp\left(-\gamma ||\mathbf{x}_i - \mathbf{x}||^2\right) \tag{3}$$

where $\gamma$ determines the width of the kernel. We use the default hyperparameters for the *svm.SVC* algorithm ($C = 1.0$ (regularization term), $\gamma = 1/[\text{n\_features} * \mathrm{Var}(\mathbf{x})]$, tolerance $=$ 0.01, cache_size $=$ 200).

The SVM was trained on ten random voxels sampled from concentric subregions at different radial distances and at three depth bins. Limiting the number to ten voxels ensured that all subregions in all participants had the same level of potential information available for decoding regardless of the size of the region. The target for the SVM was classification of the orientation of the surround during the surround-only condition. We quantified the performance of the SVM using 10-fold cross-validation and repeated the process 50 times, resampling voxels from the ROI with replacement on each iteration to yield a robust estimate of the classification performance. Consistent with (Muckli et al., 2015), the superficial layers of the target-selective region were most accurate at predicting the orientation of the surround, and we observed a significant main effect of depth on accuracy in a two-way repeated measures ANOVA in both hemispheres (left hemisphere: two-way repeated measures ANOVA $F = 13.5$, $p = 1.93 \times 10^{-4}$; right hemisphere: two-way repeated measures ANOVA $F = 19.0$, $p = 3.71 \times 10^{-5}$). We also identified a significant main effect of radius on accuracy, which appeared to peak near the border between the target and surround (left hemisphere: two-way repeated measures ANOVA $F = 3.32$, $p = 1.15 \times 10^{-2}$; right hemisphere: two-way repeated measures ANOVA $F = 3.39$, $p = 1.11 \times 10^{-2}$). However, we did not observe a consistent interaction between depth and radius in both hemispheres, with the left hemisphere showing a significant interaction (two-way repeated measures ANOVA $F = 2.70$, $p = 5.68 \times 10^{-3}$), but not in the right hemisphere (two-way repeated measures ANOVA $F = 0.465$, $p = 9.08 \times 10^{-1}$). Due to the inconsistency between hemispheres despite the symmetry of the stimulus, we did not think that we were well positioned to interpret the interactions given the relatively low statistical power afforded by our subject count. Still, it is notable that surround orientation is present at all depths in the 0-1$\sigma$ ROI, several millimeters from the surround edge.

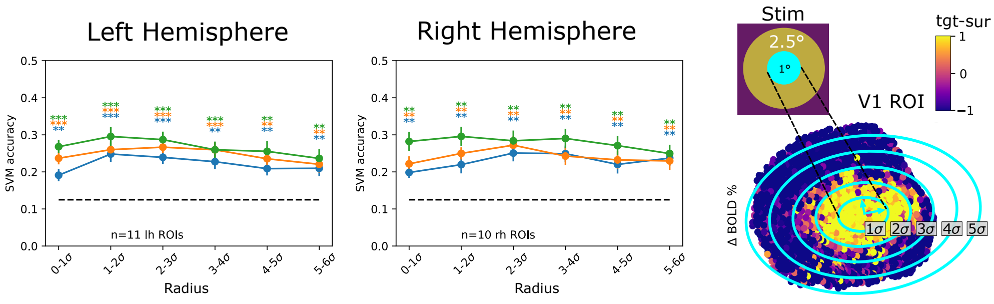


Supplemental Figure 6: **Orientation Decoding.** Classification accuracy averaged across subjects for orientation decoding analysis. Decoding analysis was performed to predict the orientation of the stimulus within the surround condition for the localizer scan. The left panel shows results for left hemisphere ROIs and the middle panel show results for right hemisphere ROIs. Color indicates depth of voxels used for decoding (deep: blue; middle: orange; superficial: green). Average accuracy is plotted at each radial bin indicated by the range on the x-axis. Sigma indicates the normalized radial coordinate determined independently for each ROI (see Methods). Error bars indicate standard error of the mean. Accuracies were compared against chance-level performance, indicated by the dotted line, using a single-sample t-test. Stars indicate statistical significance (* $p < 0.05$, ** $p < 0.01$, *** $p < 0.001$). All p-values were corrected for multiple comparisons using the Benjamini-Hochberg method (Benjamini & Hochberg, 1995). The right panel shows the concentric radial bins visualized on an example V1 patch with color indicating the localizer contrast (identical to Fig. 2E). The radial bin corresponding to the central 1-degree of the target is shown occupying the 0-1$\sigma$ bin.

Supplemental Table 26: **Left Hemisphere Depth vs Radius Orientation Classification ANOVA.** Two-way repeated measures ANOVA on orientation classification accuracy ($N = 12$; Fig. S6).

| Cases | Sphericity Corr. | Sum of Sq. | df | Mean Sq. | F | p |
|---|---|---|---|---|---|---|
| Depth | None | 0.0740 | 2.00 | 0.0370 | 13.5 | $1.93 \times 10^{-4}*$ |
| | Greenhouse-Geisser | 0.0740 | 1.09 | 0.0680 | 13.5 | $3.26 \times 10^{-3}*$ |
| | Huynh-Feldt | 0.0740 | 1.12 | 0.0660 | 13.5 | $2.97 \times 10^{-3}*$ |
| Radius | None | 0.0280 | 5.00 | 0.00600 | 3.32 | $1.15 \times 10^{-2}*$ |
| | Greenhouse-Geisser | 0.0280 | 2.55 | 0.0110 | 3.32 | $4.20 \times 10^{-2}*$ |
| | Huynh-Feldt | 0.0280 | 3.51 | 0.00800 | 3.32 | $2.52 \times 10^{-2}*$ |
| Depth * Radius | None | 0.0190 | 10.0 | 0.00200 | 2.70 | $5.68 \times 10^{-3}*$ |
| | Greenhouse-Geisser | 0.0190 | 4.88 | 0.00400 | 2.70 | $3.22 \times 10^{-2}*$ |
| | Huynh-Feldt | 0.0190 | 10.0 | 0.00200 | 2.70 | $5.68 \times 10^{-2}*$ |

Supplemental Table 27: **Right Hemisphere Depth vs Radius Orientation Classification ANOVA.** Two-way repeated measures ANOVA on orientation classification accuracy ($N$ = 12; Fig. S6).

| Cases | Sphericity Corr. | Sum of Sq. | df | Mean Sq. | F | p |
|---|---|---|---|---|---|---|
| Depth | None | 0.0560 | 2.00 | 0.0280 | 19.0 | $3.70 \times 10^{-5}*$ |
| | Greenhouse-Geisser | 0.0560 | 1.82 | 0.0310 | 19.0 | $7.37 \times 10^{-5}*$ |
| | Huynh-Feldt | 0.0560 | 2.00 | 0.0280 | 19.0 | $3.70 \times 10^{-5}*$ |
| Radius | None | 0.0510 | 5.00 | 0.00100 | 3.39 | $1.11 \times 10^{-2}*$ |
| | Greenhouse-Geisser | 0.0510 | 2.43 | 0.0210 | 3.39 | $4.45 \times 10^{-2}*$ |
| | Huynh-Feldt | 0.0510 | 3.39 | 0.0150 | 3.39 | $2.63 \times 10^{-2}*$ |
| Depth * Radius | None | 0.00600 | 10.0 | 0.000638 | 0.465 | $9.08 \times 10^{-1}$ |

Supplemental Table 28: **Orientation Decoding Statistics.** Accuracy for orientation decoding. Statistical significance is tested using one-sample two-sided t-tests with FDR-correction across all accuracy measurements using the Benjamini-Hochberg method (Benjamini & Hochberg, 1995). (Fig. S6)

| Hemisphere | Radius | Depth | Mean Accuracy | St. Dev. | N | $p$-value |
|---|---|---|---|---|---|---|
| left | $0-1\sigma$ | 0 | 0.205 | 0.0638 | 11 | $1.81\times10^{-3}*$ |
| left | $0-1\sigma$ | 0.5 | 0.234 | 0.0687 | 11 | $1.02\times10^{-3}*$ |
| left | $0-1\sigma$ | 1 | 0.266 | 0.0874 | 11 | $8.46\times10^{-4}*$ |
| left | $1-2\sigma$ | 0 | 0.224 | 0.0528 | 11 | $1.65\times10^{-3}*$ |
| left | $1-2\sigma$ | 0.5 | 0.242 | 0.0566 | 11 | $8.48\times10^{-4}*$ |
| left | $1-2\sigma$ | 1 | 0.273 | 0.0867 | 11 | $8.46\times10^{-4}*$ |
| left | $2-3\sigma$ | 0 | 0.230 | 0.0574 | 11 | $1.35\times10^{-3}*$ |
| left | $2-3\sigma$ | 0.5 | 0.249 | 0.0639 | 11 | $8.69\times10^{-4}*$ |
| left | $2-3\sigma$ | 1 | 0.271 | 0.0590 | 11 | $8.46\times10^{-4}*$ |
| left | $3-4\sigma$ | 0 | 0.210 | 0.0657 | 11 | $9.25\times10^{-4}*$ |
| left | $3-4\sigma$ | 0.5 | 0.235 | 0.0724 | 11 | $8.48\times10^{-4}*$ |
| left | $3-4\sigma$ | 1 | 0.271 | 0.0641 | 11 | $8.46\times10^{-4}*$ |
| left | $4-5\sigma$ | 0 | 0.212 | 0.0524 | 11 | $2.30\times10^{-3}*$ |
| left | $4-5\sigma$ | 0.5 | 0.229 | 0.0864 | 11 | $1.36\times10^{-3}*$ |
| left | $4-5\sigma$ | 1 | 0.252 | 0.0801 | 11 | $1.13\times10^{-3}*$ |
| left | $5-6\sigma$ | 0 | 0.224 | 0.0617 | 11 | $1.40\times10^{-3}*$ |
| left | $5-6\sigma$ | 0.5 | 0.236 | 0.0689 | 11 | $1.21\times10^{-3}*$ |
| left | $5-6\sigma$ | 1 | 0.243 | 0.0789 | 11 | $1.21\times10^{-3}*$ |
| right | $0-1\sigma$ | 0 | 0.200 | 0.0507 | 10 | $1.81\times10^{-3}*$ |
| right | $0-1\sigma$ | 0.5 | 0.213 | 0.0510 | 10 | $1.02\times10^{-3}*$ |
| right | $0-1\sigma$ | 1 | 0.250 | 0.0570 | 10 | $8.46\times10^{-4}*$ |
| right | $1-2\sigma$ | 0 | 0.222 | 0.0641 | 10 | $1.65\times10^{-3}*$ |
| right | $1-2\sigma$ | 0.5 | 0.249 | 0.0673 | 10 | $8.48\times10^{-4}*$ |
| right | $1-2\sigma$ | 1 | 0.285 | 0.0724 | 10 | $8.46\times10^{-4}*$ |
| right | $2-3\sigma$ | 0 | 0.234 | 0.0687 | 10 | $1.35\times10^{-3}*$ |
| right | $2-3\sigma$ | 0.5 | 0.254 | 0.0709 | 10 | $8.69\times10^{-4}*$ |
| right | $2-3\sigma$ | 1 | 0.284 | 0.0770 | 10 | $8.46\times10^{-4}*$ |
| right | $3-4\sigma$ | 0 | 0.235 | 0.0614 | 10 | $9.25\times10^{-4}*$ |
| right | $3-4\sigma$ | 0.5 | 0.249 | 0.0657 | 10 | $8.48\times10^{-4}*$ |
| right | $3-4\sigma$ | 1 | 0.274 | 0.0714 | 10 | $8.46\times10^{-4}*$ |
| right | $4-5\sigma$ | 0 | 0.216 | 0.0642 | 10 | $2.30\times10^{-3}*$ |
| right | $4-5\sigma$ | 0.5 | 0.232 | 0.0678 | 10 | $1.36\times10^{-3}*$ |
| right | $4-5\sigma$ | 1 | 0.260 | 0.0803 | 10 | $1.13\times10^{-3}*$ |
| right | $5-6\sigma$ | 0 | 0.226 | 0.0647 | 10 | $1.40\times10^{-3}*$ |
| right | $5-6\sigma$ | 0.5 | 0.211 | 0.0524 | 10 | $1.21\times10^{-3}*$ |
| right | $5-6\sigma$ | 1 | 0.248 | 0.0754 | 10 | $1.21\times10^{-3}*$ |

### Behavioral Results

We analyzed the behavioral responses for the orientation-discrimination task as part of a separate analysis on the relationship between cortical orientation representation and the oblique effect (Mach, 1860; Appelle, 1972). During the task, participants fixated on a small central grating stimulus that was matched in orientation to the surround. Within each trial, the stimulus was presented twice in rapid succession, after which, participants were asked to respond with a button press indicating the direction in which the orientation of the gratings had shifted. The size of the orientation shift ($\Delta\theta$) was adaptively adjusted to measure a discrimination threshold (see Methods for details).

To determine the orientation-discrimination threshold, we quantified response accuracy for each $\Delta\theta$ value within each orientation condition and fit a Weibull psychometric function (Weibull, 1951) to each subject's accuracy data using a non-linear least squares method implemented by *scipy.optimize.curve_fit* (`https://docs.scipy.org/doc/scipy/reference/generated/scipy.optimize.curve_fit.html`).

The discrimination threshold, $\Delta\theta_{th}$, was computed for a three-down, one-up staircase (79.4%). Eight of sixteen subjects had very poor performance due to $\Delta\theta$ reaching a ceiling of 16 degrees. For the remaining subjects, we noticed that many saw greatly reduced $\Delta\theta_{th}$ for the cardinal orientations over oblique orientations.

To test whether there was an effect of orientation on orientation-discrimination thresholds, we excluded subjects with poor performance by setting a criterion that they must have reached at least the discrimination threshold accuracy (79.4%) for trials with cardinal orientations (0 and 90 degrees). This yielded eight subjects that were included in the final analysis. At each orientation, we excluded those for which we were unable to estimate a $\Delta\theta_{th}$ due to poor fit quality or failure to reach threshold performance, which resulted two orientations having less than eight participants contributing data (135°: N=6 and 157.5°: N=7) . We observed a non-significant but trending effect of orientation on the orientation discrimination threshold (one-way ANOVA: $F = 2.0814$, $p = 0.0617$) (see Fig. S7). We noticed that the lowest orientation discrimination thresholds were at the cardinal orientations (0 and 90

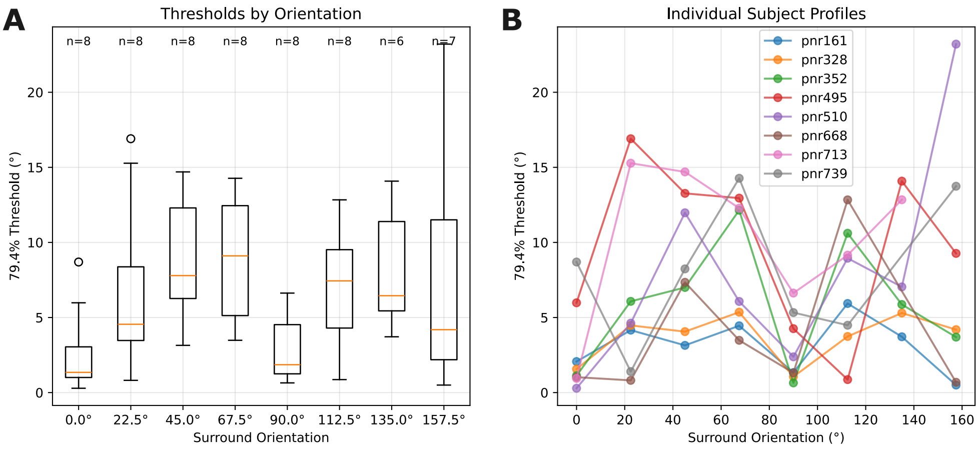


Supplemental Figure 7: **Summary of Orientation Discrimination Thresholds.** A) Orientation discrimination thresholds ($\Delta\theta_{th}$) are plotted in box and whisker plots for each orientation. Subjects that met or exceeded 79.4% accuracy on cardinal orientation trials (0 and 90 degrees) were included in the analysis. The number of subjects included within each orientation bin varied depending on the number of subjects for which a threshold could be estimated from the psychometric functions. Orange bars indicate the median $\Delta\theta_{th}$, boxes delineate the upper and lower quartiles, the bars indicate 1.5 times the interquartile range, and circles indicate outliers. B) Individual subject $\Delta\theta_{th}$ estimates are plotted as a function of orientation. Color indicates the subject profile.

degrees), indicative of the oblique effect, in which oblique orientations are not as finely resolved.

Because the adaptive staircase was applied across all conditions and was not separated based on orientation, one possibility is that the difficulty in quantifying orientation discrimination thresholds in many participants was due to the strong oblique effect. Since our estimated orientation discrimination thresholds were much lower for the cardinal orientations, it is likely that adapting $\Delta\theta$ during cardinal blocks reduced $\Delta\theta$ so that the orientations were indiscriminable during oblique blocks. Conversely, adapting $\Delta\theta$ during oblique blocks would increase $\Delta\theta$ so that accuracy became high during cardinal blocks and sensitivity was lost for estimating orientation discrimination thresholds. This would lead to inefficient sampling of orientations for some subjects and inconsistent task difficulty. Alternatively, the intersubject variability in orientation discrimination and its mean dependence on orientation condition could be due to separate effects of high intersubject variability of orientation discrimination thresholds and greater orientation discrimination thresholds at oblique angles.

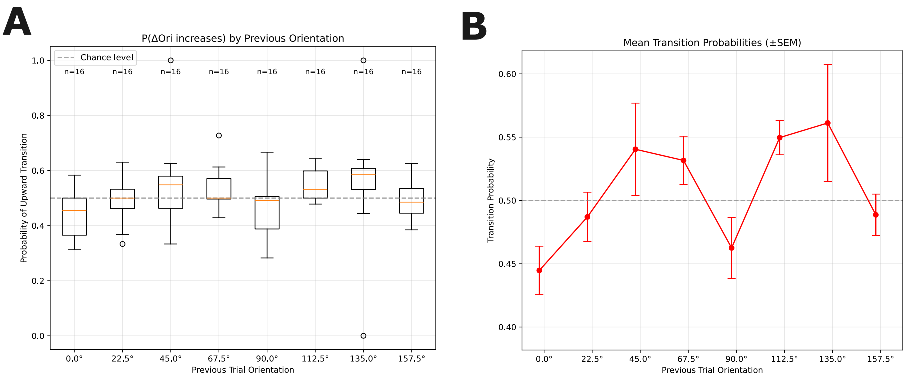


Supplemental Figure 8: **Orientation Step Transition Probabilities.** The transition probabilities for the change in orientation step ($\Delta\theta$) given the orientation on the last trial are plotted as box and whisker plots. A) Box plots show the distributions of upward transition probabilities across subjects as a function of the orientation of the trial prior to the transition. Box plots follow the same conventions as Fig. S7. B) The mean upward transition probabilities across subjects are plotted against the orientation of the trial prior to the transition. Error bars indicate standard error of the mean.

To more directly test the effect of orientation discrimination of task difficulty, we computed the direction of transitions (upward or downward) of $\Delta\theta$ between trials. We reasoned that task difficulty would be reflected in the probability of transitioning to larger or smaller $\Delta\theta$ within orientation conditions. A participant performing at equilibrium will observe an equal probability of transitioning to a higher or lower $\Delta\theta$. If the probability of transitions varied significantly from equilibrium between orientation conditions, this would suggest varying task difficulty across conditions. We quantified the probability of the transition given the orientation of the previous trial by considering all trials in which upward or downward $\Delta\theta$ transitions occurred. We found that orientation had a significant effect on the upward and downward transition probabilities (one-way ANOVA: $F = 2.4953$, $p = 0.0199$) (see Fig. S8) suggesting that participants were not performing at equilibrium during all orientation conditions. Upward transitions were more likely during oblique trials and downward transitions were more likely during cardinal trials, suggesting that cardinal trials systematically shifted the $\Delta\theta$ downward while oblique trials systematically shifted the $\Delta\theta$ upward. This confirms our suspicion that task difficulty varied significantly by orientation condition.

As mentioned in the methods, the behavioral task was irrelevant to the contextual modulation paradigm and was only useful insofar as it encouraged participant engagement. However, because the behavioral results were confounded by the oblique effect, making the task too difficult for many participants, we could not use task performance as a reliable measure of participant engagement.

### Inclusion Criteria

Inclusion criteria were applied to each ROI separately. Details on inclusion criteria are described in Materials and Methods. Table S29 reports each of the original ROIs from 16 participants and whether they were included for analysis.

Supplemental Table 29: **ROI Inclusion/Exclusion.** V1 target-selective ROIs were evaluated separately for each subject in each hemisphere and independently subject to inclusion criteria. The reason for exclusion is listed with excluded ROIs. 21 ROIs in 12 participants were included.

| **Participant ID** | **Hemisphere** | **Inclusion/Exclusion** |
|---|---|---|
| pnr102 | lh | Excluded; low SNR and no discernible target-selective region |
| pnr102 | rh | Excluded; low SNR and no discernible target-selective region |
| pnr143 | lh | Included |
| pnr143 | rh | Excluded; poor boundary fit |
| pnr161 | lh | Excluded; inadequate number of significantly target-selective voxels ($p < 0.01$) |
| pnr161 | rh | Excluded; poor boundary fit |
| pnr256 | lh | Included |
| pnr256 | rh | Included |
| pnr328 | lh | Included |
| pnr328 | rh | Included |
| pnr352 | lh | Excluded; no discernible target-selective region |

| Participant ID | Hemisphere | Inclusion/Exclusion |
|---|---|---|
| pnr352 | rh | Excluded; inadequate number of significantly target-selective voxels ($p < 0.01$) |
| pnr495 | lh | Included |
| pnr495 | rh | Included |
| pnr510 | lh | Included |
| pnr510 | rh | Included |
| pnr579 | lh | Excluded; poor boundary fit |
| pnr579 | rh | Included |
| pnr668 | lh | Included |
| pnr668 | rh | Excluded; poor boundary fit |
| pnr685 | lh | Included |
| pnr685 | rh | Included |
| pnr713 | lh | Included |
| pnr713 | rh | Included |
| pnr739 | lh | Included |
| pnr739 | rh | Included |
| pnr756 | lh | Included |
| pnr756 | rh | Included |
| pnr822 | lh | Included |
| pnr822 | rh | Included |
| pnr947 | lh | Excluded; no discernible target-selective region |
| pnr947 | rh | Excluded; no discernible target-selective region |
